# Functional traits predict species responses to environmental variation in a California grassland annual plant community

**DOI:** 10.1101/2020.11.17.387076

**Authors:** Gaurav S. Kandlikar, Andew R. Kleinhesselink, Nathan J.B. Kraft

## Abstract

1. Turnover in species composition and community-wide functional traits across environmental gradients is a ubiquitous pattern in ecology, and is generally assumed to reflect shifts in trait optima across these gradients. However, the demographic processes that give rise to these trait turnover patterns at the community level remain unclear.
2. We asked whether shifts in the community-weighted means of three key functional traits across an environmental gradient in a southern California grassland reflect variation in the trait-performance relationship across the landscape.
3. We planted seeds of 17 annual plant species in cleared patches with no competitors, and quantified the lifetime seed production of 1360 individuals. We then asked whether models that included trait-environment interactions help explain interspecific variation in demographic responses to the environment. This allowed us to evaluate whether observed shifts in community-weighted mean traits matched the direction of any trait-environment interactions detected in the plant performance experiment.
4. Our results indicate that commonly-measured plant functional traits help explain variation in species responses to the environment – for example, high-SLA species had a demographic advantage in soils with high soil Ca:Mg levels, while low-SLA species had an advantage in low Ca:Mg soils. We also found that shifts in community-weighted mean traits often reflect the direction of these trait-environment interactions, though not all trait-environment relationships at the community level reflect interactive effects of traits and environment on species performance.
5. Our results support the value of plant functional traits for predicting species responses to environmental variation, and highlight a need for more detailed evaluation of how trait-performance relationships change across environments to improve such predictions.

## Introduction

Understanding how environmental variation shapes the diversity and dynamics of plant communities is a fundamental challenge in ecology. In addition to variation in species composition (Whittaker 1960; Janzen 1967), turnover in the functional traits of plant communities across abiotic gradients has emerged as a ubiquitous pattern across ecosystems (Cavender-Bares et al. 2004; Hulshof et al. 2013; Bjorkman et al. 2018; Jardine et al. 2020). These functional traits reflect key physiological and life history strategies of plants, which ultimately determine variation in plant fitness across different environments (Violle et al. 2007; Reich 2014). One of the most common ways for plant ecologists to study trait-environmental relationships has been to quantify variation in community-weighted mean (CWM) of functional traits across landscapes. CWM trait values are calculated as species’ trait values weighted by their relative biomass or cover, and reflect the functional properties of the dominant plant species growing in a community (Grime 1998; Garnier et al. 2004). Across ecosystems, communities with less harsh abiotic conditions (e.g. lower drought stress, higher resource availability) tend to be dominated by plants with functional traits that generally reflect resource-acquisitive strategies (e.g. higher specific leaf area or leaf N concentrations (Reich 2014)), and vice-versa in environments that are less favorable for plant growth. Shifts in CWM traits are often assumed to reflect variation in trait optima across environmental gradients, with species whose traits closely match CWM expected to have highest fitness (Ackerly 2003; Shipley et al. 2006; Enquist et al. 2015).

Although shifts in the functional traits of plant communities across environmental gradients is well-documented, the demographic processes driving these pattern remain unclear (Salguero-Gomez et al. 2018; Laughlin et al. 2020). As a result, predicting how variation in species functional traits drives variation in community composition – one of the key promises of functional trait ecology (McGill et al. 2006; Westoby and Wright 2006)– remains a challenge. For example, Muscarella and Uriarte (2016) found that a substantial portion of tree species in a tropical forest were more abundant in sites where their traits were more dissimilar from the site’s CWM, contrary to predictions of the hypothesis that CWM shifts reflect shifts in trait optima. Part of the challenge is that we lack a clear understanding of whether CWM trait shifts reflect variation in the relationship between functional traits and the vital rates (e.g. germination rate, fecundity, survival rate) that ultimately determine species performance across landscapes (Shipley et al. 2016).

One path forward is to pair observed shifts in CWM traits with analyses that evaluate the interactive effect of traits environments on species’ demographic rates (Laughlin and Messier 2015; Swenson et al. 2020). For example, in one of the few studies that has investigated whether CWM trait shifts reflect variation in trait optima, Laughlin et al. (2018) found CWM shifts in leaf, root, and reproductive functional to be unreliable predictors of how traits influence survival rates of plants in a pine forest system, also contradicting the predictions of the idea that CWM trait shifts reflect shifting trait optima. It is important for such analyses to quantify species fitness based on their vital rates or population growth rates rather than species abundance measured at a single time point, which can be influenced multiple abiotic and biotic processes (e.g. dispersal, competition, natural enemies) and is thus a poor proxy for intrinsic fitness (e.g. Fox 2012; McGill 2012).

The impact of changing trait-performance relationships on CWM traits can be expected to take one of a number of forms, some of which are illustrated in Fig. 1. If trait-performance relations remain constant across an environmental gradient (Fig. 1B), any observed CWM trait shifts likely reflect the effects of species interactions or other processes rather than shifting trait optima in terms of species’ intrinsic responses to the landscape. Trait-performance relationships may differ in magnitude but not in sign across a gradient in a way that matches observed shifts in CWM traits (Fig. 1C). Such trait-performance relationships with the same sign across the environmental gradient would not by themselves result in differential distribution of traits across the landscape, but provide weak support that CWM trait shifts reflect shifting trait optima. The strongest evidence that CWM trait shifts reflect shifting trait optima would be if the sign of the trait-performance relationship changes across the gradient in a way that is consistent with the CWM trait patterns (Fig. 1D). It is also possible that we find strong trait-environment interactions when looking at the vital rates even when there are no observed CWM trait shifts. This might indicate that other processes obscure underlying trait-performance relationships. A major challenge in testing for concordance between CWM trait shifts and variation in trait-performance relationships has been that quantifying how trait variation influences species demography across landscapes is very data-intensive, requiring plant performance data across large temporal and spatial gradients. Although such data are becoming increasingly available for a wide range of perennial plants (e.g. Salguero-Gomez et al. 2014), short-lived plant communities, where we can quantify relevant vital rates on fairly short time scales, offer an ideal system in which to test for concordance between trait-performance relationships and CWM trait shifts.

**Figure 1:**
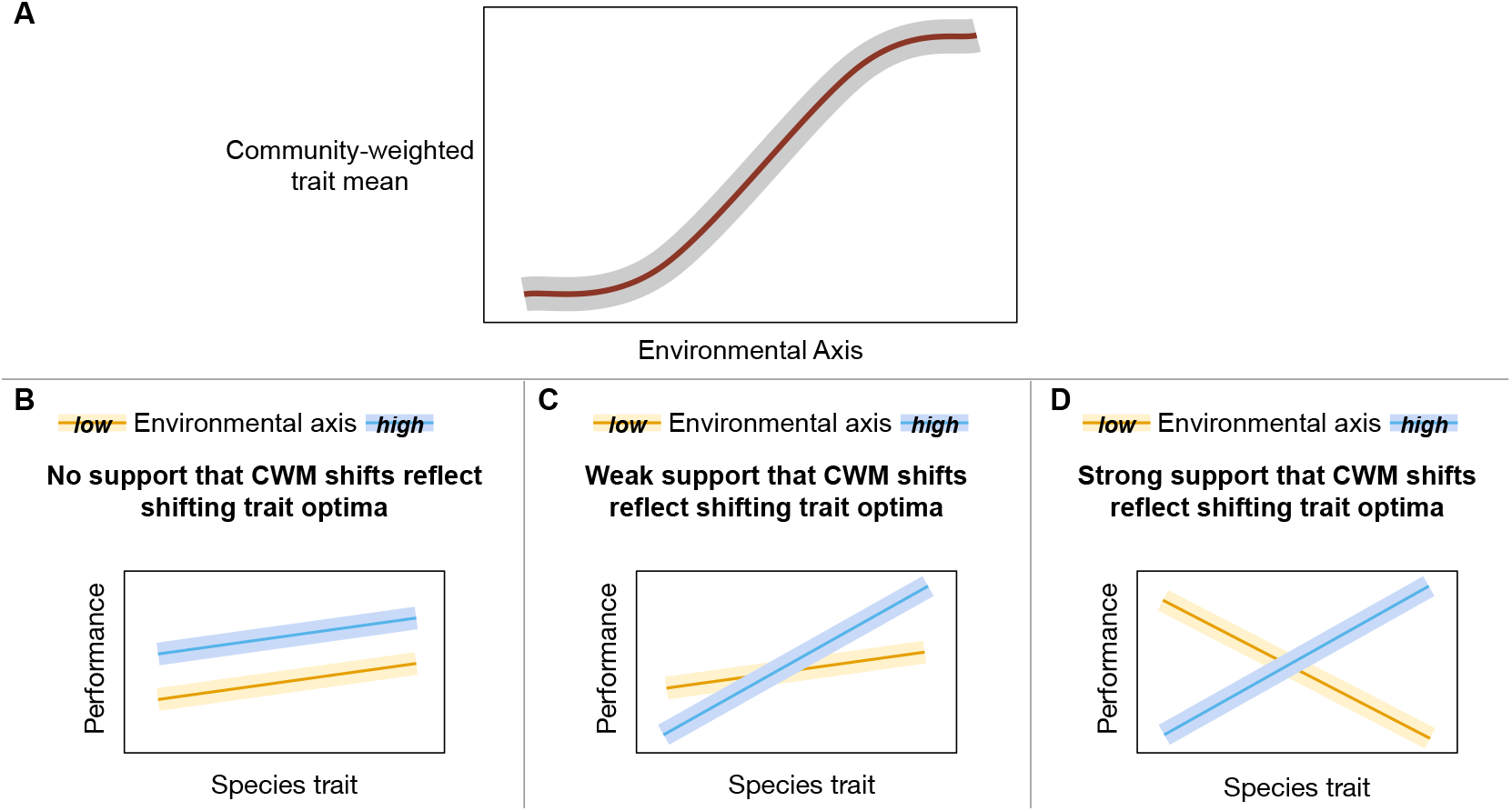
A) Variation in community-weighted mean (CWM) functional traits across gradients is a common pattern in plant communities, though whether or not such variation in CWM traits reflects shifts in trait optima across environmental gradients. Here we evaluate whether CWM shifts in plant functional traits reflect shifts in trait-performance relationships across key environmental gradients. Panels B-D illustrate how trait-performance relationships might vary across environments. B) The trait-performance relationship may be identical at opposite ends of the environmental gradient, indicating that other factors (e.g. dispersal limitation) might drive observed shifts in CWM traits. We interpret this as a lack of evidence that CWM trait-environment relationships reflect variation in trait optima across the environment. C) The trait-performance relationship may change across the environmental gradient in a direction that is consistent with observed CWM shifts, but the sign of the trait-performance relationship may be the same at either end of the gradient. We interpret this as providing weak evidence that CWM shifts reflect changing trait optima. D) The sign of the trait-performance relationship may change across the gradient, such that species with low trait values have a relative advantage at the low end of the environmental gradient, and vice versa at the high end of the gradient. We interpret this as strong evidence that CWM shifts reflect changing trait optima.

**Figure 2:**
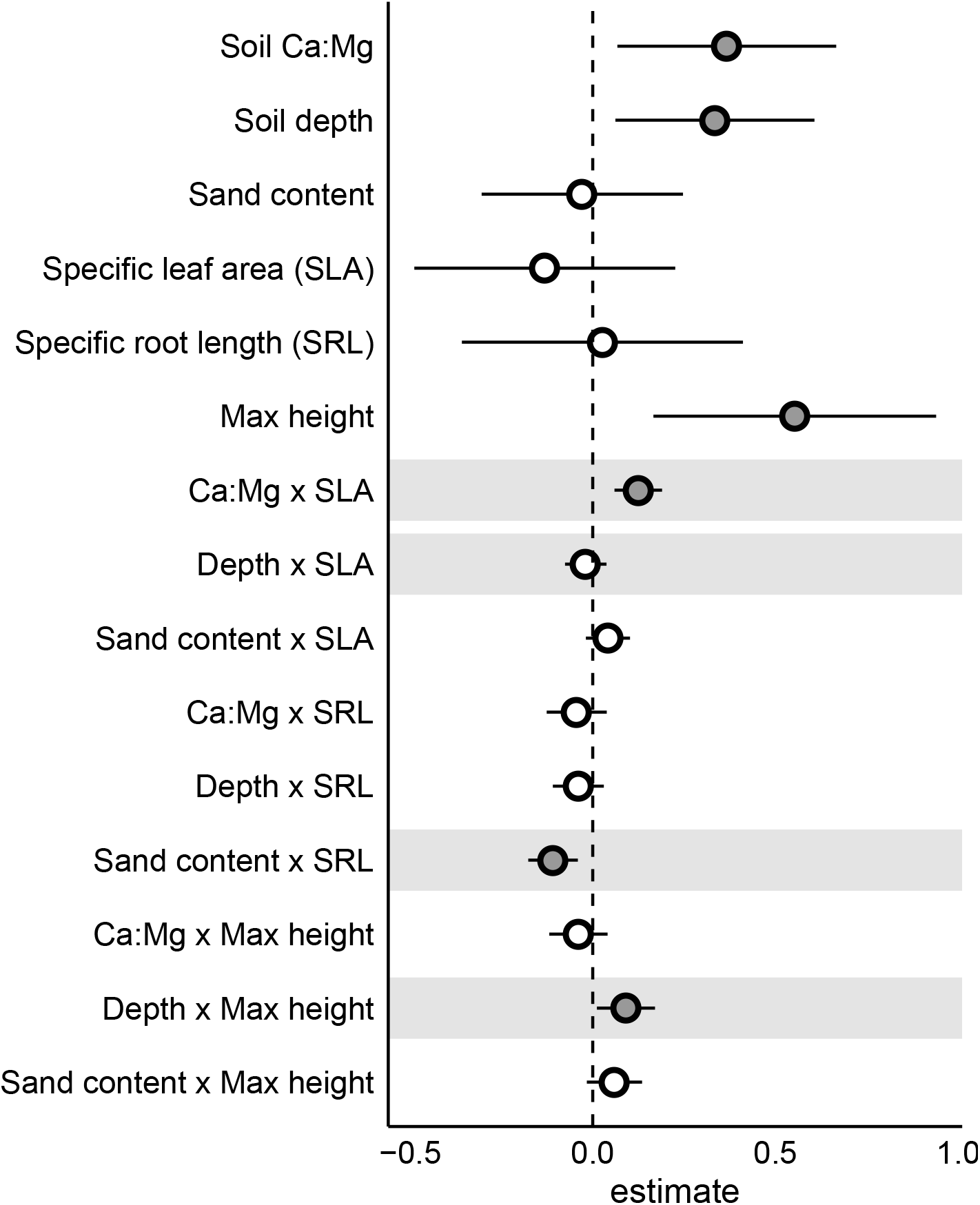
Standardized effects of environmental variables, functional traits, and their interactions on seed production. Grey points indicate those effects whose 95% confidence intervals (indicated by thin bars) do not overlap zero. Horizontal grey bars indicate the four significant trait-environment relations we observed in the CWM trait analysis.

In order to address this longstanding question of how whether trait shifts at the community level reflect variation in trait optima across the landscape, here we compared CWM trait shifts to the interactive effects of traits and environment on species’ fitness in a southern California serpentine annual grassland community. We surveyed the plant community at sites that captured a wide range of variation in soil Ca:Mg, sand content, and depth – three axes of abiotic variation that are known to be important in such serpentine communities. To capture various dimensions of plant ecological strategies, we quantified community-wide variation in one leaf trait (specific leaf area), one root trait (specific root length), and one whole-plant trait (maximum height). In a parallel experiment, we quantified the intrinsic fitness (lifetime fecundity of individuals growing without competitors) of 17 annual plant species that naturally occur in this community and that capture a wide range of functional variation. We then asked whether observed CWM trait shifts reflect trait-environment interactions that shape variation in species’ fecundity across this gradient. Our results show that shifts in CWM traits can provide valuable information into how trait optima shift across gradients, but also caution against predicting species responses to environmental variation on the basis of shifts in CWM traits alone.

## Methods

### Study system

We studied trait-environment relations in the grassland community at the University of California Sedgwick Reserve in southern coastal California. This region experiences a Mediterranean climate of cool, wet winters and long, dry summers. Plant phenology in this system is driven largely by the rainfall regime: seeds of plants germinate with early-season rain storms in November and December, and plants begin to reproduce and senesce with the onset of summer droughts (though there is substantial variation in the timing of reproduction among species (Godoy and Levine 2014; Kraft et al. 2015)). The reserve encompasses significant topographic and edaphic heterogeneity, including oak-savanna, coastal sage scrub, and California grassland communities. Our study focused on a part of the reserve with serpentine-derived soils that are dominated by invasive *Avena* and *Bromus* spp. In this area, rocky serpentine outcrops (“hummocks”) are embedded within a matrix of deep, clay soil. The outcrops are considerably less vegetated than the matrix soils, and act as spatial refuges for several native plant species (Gram et al. 2004). We studied trait-environment interactions at 16 sites on this landscape, with 10 sites located on serpentine hummocks and 6 in the matrix.

### Quantifying species performance across the landscape

In November 2015, before the first major rain storm of the season, we cleared all existing vegetation in 2m X 3m plots at each of our 16 focal sites. At each site, we sowed five replicate plots with the equivalent of 20-60 viable seeds each of our 17 focal species (Table S2) on a grid with 15 cm spacing between each species. We collected seeds from hundreds of plants growing across Sedgwick Reserve in the spring prior to this study, and seeds were homogenized among sources before planting to ensure that local adaptation (Rajakaruna and Bohm 1999) or maternal effects (Germain and Gilbert 2014) did not systematically drive variation in plant performance across sites in our experiment.

In February 2016, we counted the number of germinants of each focal species in our experimental plots, and thinned each plot to leave only two individuals of each focal species. In March, we further thinned each point down to a single individual of each species, and weeded around this focal individual to ensure that it was not competing with other plants in a 15cm radius. Between April-June 2016, we quantified the total seed output of each focal individual in our experiment, for a total of 1360 individuals (17 species * 16 sites * 5 plots per site) tracked across the environment (see Appendix S1 for details on how we estimated total seed output). This design let us quantify the germination rate and the per-germinant seed production (fecundity) in the absence of competitors for each species at each site. Both of these vital rates are known to be important determinant of annual plant demography in this community (Levine and HilleRisLambers 2009), but we focus only on fecundity as a measure of species performance in the remainder of this study because the functional traits we measured most clearly relate to the growth of plants after germination.

### Measuring compositional turnover across the landscape

In Spring 2017 (the year after the experimental assessment of plant performance), we surveyed five undisturbed plots (0.5×0.5m) adjacent to the experimental plots at each of our 16 sites to characterize the vascular plant community composition. These community survey plots were spaced evenly on a 10m transect located alongside the cleared plots in which we had experimentally quantified plant performance in the prior year. In each plot, we visually estimated the total (absolute) cover of each of species in early April, and again in early June.

### Functional trait measurement

Kraft et al. (2015) had previously measured 11 functional traits (Table S1) that to capture ecologically important variation in leaf, root, whole-plant, and reproductive functioning of plant species for most species in our demography experiment (12 out of 17 species; note that *U. lindelyi* was misidentified *Agoseris heterophylla* in that study). In Spring 2016, we measured the same set of traits for the five species in our experiment that were not part of Kraft et al. (2015)’s study: *Bromus madritensis*, *Chaenactis glabriuscula*, *Hordeum murinum*, *Micropus californica*, and *Vulpia microstachys*. We followed the protocols detailed in Kraft et al. (2015) to measure traits from 5-8 individuals growing in 0.7*0.7m plots at three of the matrix sites in our experiment, which we had sowed with seeds of all 17 annual plant species for a total sowing density of 8g viable seeds/m^2^. In Spring 2017, we measured the same set of functional traits on the 38 of the most common annual plant species encountered within the community composition plots (of the species for which we could not measure at least one of the focal traits, mean cover of these species in sites where they were present was < 5%).

Our analysis focuses on three traits that capture distinct dimensions of plant ecological strategies and that were largely uncorrelated in a principal components analysis of the traits we measured (Fig. S1): specific leaf area (SLA), specific root length (SRL), and maximum height. SLA, the ratio of leaf area to dry mass, is strongly linked to species’ position along the leaf economics spectrum (Wright et al. 2004) and at a global scale is positively correlated to photosynthesis and growth rate (Adler et al. 2014). SRL, the ratio of fine root length to dry mass, reflects the area over which roots can uptake resources relative to biomass investment, is an important component of the root economics spectrum (Laliberte 2016; Weemstra et al. 2020). At both a global scale (Weemstra et al. 2016) and within our study (Fig. S1), SRL is largely uncorrelated with SLA. Species with higher SRL tend to have superior nutrient acquisition capability, but are generally more susceptible to attack by pathogenic microbes (Eissenstat 1992; Laliberte et al. 2014). Maximum height is a globally relevant trait (Díaz et al. 2016) that integrates across various dimensions of ecological strategy and can indicate the ability of adult plants to preempt and intercept light (Westoby et al. 2002). The 17 focal species of our performance experiment reflected a wide range of variation observed across the plant community for these three traits (Fig. S2)

### Environmental sampling

We quantified various soil chemical and physical characteristics to identify the primary axes of environmental variation among our study sites. We measured gravimetric water content ((weight of fresh soil - weight of dry soil)/weight of dry soil) in the early- and mid-growing season (March and April, respectively), and summarized across these measurements to estimate the average soil moisture at each site. At each site, we also collected soil for analysis by A&L Western Agricultural Laboratories (Modesto, CA) for a variety of soil chemical and physical properties: soil organic matter, P (Weak Bray and Olsen methods), K (ppm), Mg (ppm), Ca (ppm), Na(ppm), pH, CEC, NO_3_, SO_4_, NH_4_, and soil texture (sand, silt, and clay content). We collected the soil for these analysis from three points arranged in between the five experimental plots, and homogenized within site prior to analysis. We also programmed iButtons (Maxim Integrated) to log temperature at 2-hr intervals, and used these data to quantify the average daily maximum temperature at each site. To avoid direct solar radiation on iButtons, we placed them in anchored PVC tubes with holes for airflow. Based on a PCA of all environmental variables (Fig. S3), we focus on soil Ca:Mg, soil sand content, and soil depth as biologically relevant and largely uncorrelated environmental variables that capture the primary axes of abiotic variation among our study sites.

### Analysis

#### Quantifying community-weighted trait turnover across the landscape

We used the community composition and functional trait measurements to calculate the community-weighted mean (CWM) trait values, which represent the mean trait value of all species growing at a site, weighted by the species’ relative cover. We calculated the CWM for each trait (*t*) at each of our 16 sites (*s*) by averaging across the CWM of the five plots *p* at each site as follows: 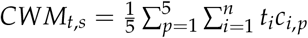, where *n* is the number of species found in each plot, *t_i_* is the mean trait value of species *i*, and *c_i_*_,*p*_ is the relative cover of species *i* in the plot *p*. We then evaluated whether CWM traits vary across the environmental gradient in our study with simple bivariate linear regressions between each of the three focal traits and each of the three focal environmental characteristics (*α* = 0.05). We also tested for evidence of nonlinear trait-environment relations by including a quadratic term in the predictor (environmental) variables.

#### Quantifying the functional trait basis for variation in species performance across the environment

We next asked whether observed CWM trait shifts across the environmental gradients in our study reflect variation in the trait-performance relationship in the demography experiment. We used `glmmTMB` (Brooks et al. 2017) to fit a generalized linear mixed effects model with each focal individual’s seed production as the response variable, and with the three focal environmental variables, three focal functional traits, and nine pairwise trait-environment interaction terms as predictors. The model also included random effects for species identity and site, and was fit with a zero-inflated negative binomial error structure. We log-transformed all functional trait values, and scaled all parameters to help with model convergence. We used `performance` (Lüdecke et al. 2020) to quantify model fit and to ensure that the model did not suffer from colinearity of predictors (variation inflation factors < 2), and used `DHARMa` (Hartig 2020) to evaluate the model residuals. We also compared the AICc of the model with all trait-environment interaction terms to a null model and one with only the main trait and environment effects as predictors to verify that the interaction terms were supported (ΔAICc > 2 in favor of the model with trait-environment interactions). Finally, we used `effects` (Fox and Weisberg 2018) to evaluate the trait-performance relationship at the highest and lowest value of the environmental gradient in our study based on the marginal effects of the model. We considered trait-environment interactions that were significant in the model, but whose slope did not change sign across the environmental gradient, as weak evidence that CWM trait shifts reflect shifts in trait optima across the landscape (Fig. 1C). If the sign of the trait-performance sign shifted in the direction predicted by CWM trait shifts, we considered this as strong evidence that CWM trait shifts reflect shifts in trait optima across the landscape (Fig. 1D).

We conducted all analyses in R v. 3.6.3 (R Core Team 2020) and provide code to recreate all analyses in appendix S2. All data are available as supplementary files and will be deposited in to an archival repository prior to publication.

## Results

### Community-wide trait turnover at Sedgwick

The plant species in our study system vary considerably in their functional traits. Across the 55 species we observed across the landscape, there was 3 fold variation in SLA (5th percentile = 124.83*cm*^2^/*g*, 95th percentile = 433.8*cm*^2^/*g*), 9 fold variation in SRL (5th percentile = 32.26*m*/*g*, 95th percentile = 290.67*m*/*g*), and 10 fold variation in Maximum Height (5th percentile = 11.38cm, 95th percentile = 108.7cm). This trait variation was strongly structured along various environmental axes in our study. We observed strong positive relationships between CWM SLA and soil Ca:Mg and soil depth (Fig. 3A-B), a strong negative relationship between CWM SRL and soil sand content (Fig. 3C), and a strong positive relationship between CWM max height and soil depth (Fig. 3D). We also found evidence that CWM SRL tends to be highest at intermediate values of Ca:Mg (Table S3).

**Figure 3:**
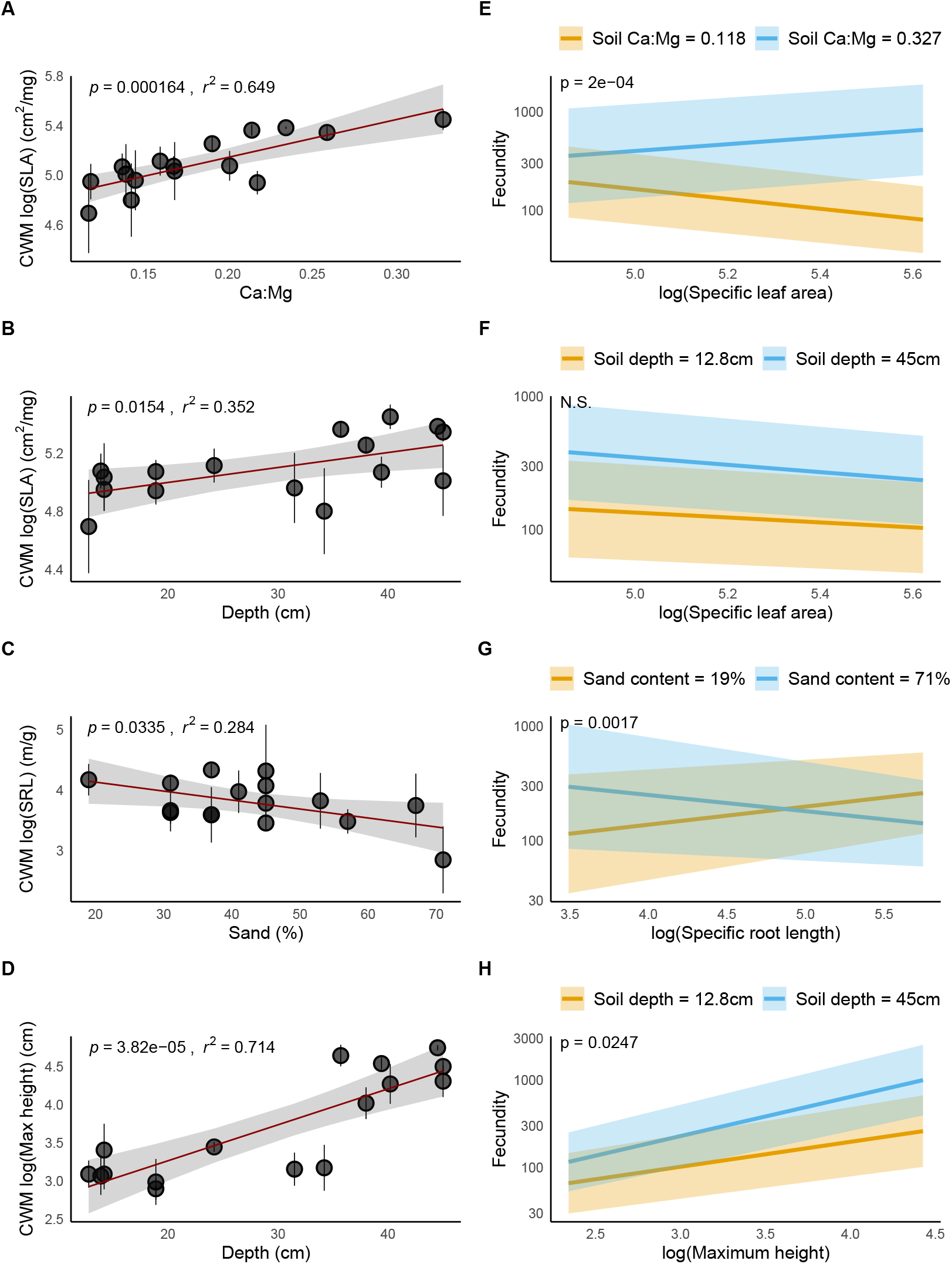
Plots A-D show the four significant relationships we observed between environmental variables and community weighted mean traits (points indicate means and bars show the 95% confidence interval at each site). Panels E-H show the corresponding trait-environment interactions from our GLMM, with yellow lines showing the trait-performance relationship at the lowest value of the environmental variable observed in our study, and the blue lines at the highest. Translucent bands in all panels represent the 95% confidence intervals. Note that model predictions were made with scaled values of trait and environmental predictors, which were back-transformed onto their original scale for visualization. Plots of all non-significant trait-environment relationships are available in Fig. S4, and 3D surfaces for each interaction in Fig. S5.

### Environmental and functional trait drivers of variation in species performance

The fixed effects of our GLMM with all the main and interactive effects explained 18% of the variation in seed production (Marginal R^2^ = 0.18), and the random effects of species and site explained an additional 19% of the variation (Conditional R^2^ = 0.37). The model included significant positive main effects of soil Ca:Mg (p = 0.016) and soil depth (p = 0.016), indicating that seed production was higher in sites with higher Ca:Mg and deeper soils, irrespective of plant traits (Fig 2). The main effect of maximum height was also significant and positive (p = 0.005), indicating higher seed output from larger-statured plants across the environmental gradient (Fig 2).

The model also provided evidence that three out of the four significant relationships between CWM traits and environmental variables reflect variation in trait-performance relations across the environmental gradient. We found strong evidence that the positive relationship between CWM SLA and soil Ca:Mg (Fig. 3A) reflects a shift in the trait-performance relationship on this landscape. Seed production was shaped by a significant positive interaction between SLA and soil Ca:Mg, which caused the sign of the SLA-performance relationship to shift from negative in the lowest-Ca:Mg site to positive in the highest Ca:Mg site (Fig 3E). We found similarly strong evidence that the negative relationship between CWM-SRL and soil sand content (Fig. 3C) reflects shifts in trait-performance relationships. Seed production was shaped by a significant negative interaction between SRL and soil sand content, which caused the sign of the SRL-performance to shift from positive in the least sandy site, to negative in the most sandy sites in our study (Fig 3G). The model also provided weak evidence that the positive relationship between CWM max height and soil depth (Fig. 3D) reflects shifts in trait-performance relationships. Seed production was shaped by a significant positive interaction between maximum height and soil depth, though this interaction only reflected a more positive max height-performance relationship in deeper than in shallower soils, rather than a change in the sign of the trait-performance relationship across the soil depth gradient (Fig. 3H). The model did not provide any evidence that the negative CWM SLA-soil depth relationship (Fig. 3B) reflected a change in the SLA-performance relationship across the soil depth gradient in our study (Fig. 3F). Finally, in no case did the model identify a trait-environment interaction in the demography experiment for which we did not also find a corresponding relationship in the CWM trait analysis (Fig. S4).

## Discussion

Turnover in community-weighted trait means across environmental gradients is a ubiquitous pattern in nature, but whether these patterns reflect shifts in trait-performance relations across environmental gradients remains poorly understood (Shipley et al. 2016). As a result, predicting plant species’ demographic responses to environmental variation on the basis of their functional traits remains challenging (Laughlin and Messier 2015). Quantifying trait-performance relations across environmental gradients at the community level is a key step in improving our ability to project how plant communities will respond to environmental change. Here, we asked whether patterns of community-weighted mean trait turnover in three key functional traits reflect variation in the trait-performance relationships across three abiotic gradients in a southern California serpentine grassland community. We found evidence that three out of the four significant trait-environment relationships at the community level reflect shifts in trait-performance relationships across the gradients. Quantifying how traits mediate species’ demographic responses across their life remains a key step in improving our ability to use functional traits to predict plant community responses to environmental variation.

We found three trait-environment interactions structuring both the whole plant community as well as variation in the seed production of focal species in our experiment on this landscape. The positive relationships between CWM SLA and soil Ca:Mg is consistent with other studies that have found lower CWM SLA in sepentine soils than in soils with weaker serpentine characteristics (Fernandez-Going et al. 2013), and with the general finding of increasing CWM SLA in more fertile soils (e.g. Cornwell and Ackerly 2009). Our GLMM analysis of the focal species’ seed production suggests that this community-level pattern may arise in part because the value of SLA that confers the optimal fitness (measured here as the intrinsic fecundity of plants when not facing competitors) shifts along the Ca:Mg gradient. Our model included a significant positive main effect of soil Ca:Mg, indicating that all plants performed better in high-Ca:Mg than low-Ca:Mg soils, irrespective of their traits. However, the magnitude SLA-Ca:Mg interaction term indicates that lower SLA was associated with higher intrinsic fitness at soils with low-Ca:Mg, and vice-versa in high-Ca:Mg soils (Fig. 3E). This is consistent with the general expectation of lower SLA being correlated with a suite of traits that confer plants greater tolerance of abiotic stress, at the cost of a relative disadvantage when abiotic stress is less limiting (Wright et al. 2004; Sterck et al. 2006). For example, lower SLA was correlated with higher water use efficiency (Fig. S1), which may give low-SLA species a relative advantage in low-Ca:Mg sites, which also tended to have lower soil moisture in this system (Fig. S3).

We also found a negative relationship between CWM SRL and sand content (Fig. 3C). This relationship is contrary to Laughlin et al. (2018), who found a positive CWM SRL-soil sand content relationship and a positive interactive effect of SRL and soil sand content on plant survival in a pine-dominated forest in Arizona. This discrepancy may have arisen in part because in our system, soil sand content was generally much higher on serpentine hummocks that were also characterized by low soil moisture and organic matter (Fig. S3). In this context, the negative relationship between CWM SRL and sand content is consistent with the more general expectation of low SRL indicating a resource-conservative strategy that allows plants outperform species with resource-acquisitive strategies in more stressful conditions (Reich 2014). Moreover, our analysis of trait and environmental predictors of seed production provides strong evidence that this community-level pattern is in part driven by low-SLA species having higher intrinsic fecundity in sandy soils, and vice-versa in soils with low sand content (Fig. 3C). Understanding the drivers of variation in trait-environment interactions among different plant communities remains a key challenge in building towards a more predictive trait ecology (Funk et al. 2016), and may be achieved with more studies that couple observational studies at the community level with species-level analyses of trait-environment interactions.

The third trait-environment interaction for which we found evidence for in both the observational study and in our demography experiment was the positive interaction between soil depth and maximum height (Fig. 3D). This finding of a positive CWM max height-soil depth relationship is consistent with the distribution of plant height along soil depth gradients in other Mediterranean grassland communities (Bernard-Verdier et al. 2012). This community-level pattern may be driven by a positive interactive effect of maximum height and soil depth on intrinsic fecundity (Fig. 3H). However, this interaction term provides only weak evidence that the turnover in CWM max height across soil depth reflects a shifting trait-performance relationships, because seed production was also influenced by a significant positive main effect of species’ max height. In other words, even though the relative advantage to taller species is diminished in shallower vs. deeper soils, tall species had higher intrinsic fecundity than short species across the depth gradient. Although the interactive effect of maximum height and soil depth on intrinsic fecundity alone may not be sufficient to drive trait shifts across the landscape, trait-performance relationships in other vital rates may compound this effect to give rise the community-wide trait turnover in maximum height. For example, Kraft et al. (2015) previously found that taller species have a competitive advantage over shorter species in a pairwise competition experiment conducted on matrix soils in this landscape. Shallower soils on serpentine hummocks are also characterized by lower density of vegetation (Gram et al. 2004) and potentially less severe light competition, which could provide a competitive advantage to shorter species if there is a tradeoff in aboveground vs. belowground competitive ability (DeMalach et al. 2016). Future studies that investigate trait-performance relationships in various demographic processes will be critical for understanding how plant traits determine overall population growth rates and this influence community assembly processes across landscapes.

Although three three of the four CWM trait-environment correlations in this study seem to at least qualitatively reflect the direction of the trait-environment interaction in terms of species’ intrinsic fecundity, we did not find such evidence for the positive CWM SLA-soil depth correlation (Fig. 3B,F). This raises the question of what might drive the community-level association between species’ SLA and soil depth. It is possible that the rather than influencing how intrinsic fecundity of species responds to variation in soil depth, SLA might instead mediate the response of other vital rates to this environmental gradient. It is also possible that the CWM SLA-depth relationship arises due to trait-environment interactions between other correlated traits or environmental variables which are not reflected in our analysis of intrinsic fecundity (Marks and Lechowicz 2006; Laughlin and Messier 2015). In general, that the CWM SLA is strongly correlated with soil depth even though SLA does not appear to mediate variation in intrinsic fecundity across the soil depth gradient highlights the potential pitfalls in predicting species’ demographic responses to environmental gradients on the basis of community-wide patterns of trait turnover.

Our analysis also allows us to ask whether any trait-environment interactions mediate variation in species performance but do not appear to turn over across the environmental gradient at the community level. We did not find any evidence for trait-environment interactions influencing species performance that did not manifest in CWM trait turnover (Fig. S4). This suggests that in our annual grassland system, trait-environment interactions that shape variation in species’ intrinsic fecundity in different environments do manifest in trait turnover at the community level. In comparison, Laughlin et al. (2018) found a strong negative interactive effect of SLA and soil C:N on plant survival, but did not observe a negative CWM SLA-C:N relationship at the community level. It is possible that community patterns in annual-dominated communities are more sensitive to underlying species-level trait-environment interactions than the perennial system in Laughlin et al. (2018)’s study. Coupling empirical studies with simulations of community dynamics in systems dominated by plants of differing life histories will help to build a more general understanding of the mapping between species- and community-level trait-environment interactions.

Our overall finding that shifts in CWM traits across environmental gradients often reflect shifts in trait-performance relations but are not perfect predictors of trait optima is consistent with other studies that have investigated variation in CWM traits and species performance on a landscape (Muscarella and Uriarte 2016; Laughlin et al. 2018). However, our study also had some important limitations. First, the 17 species we used in our experiment to quantify intrinsic fecundity across the landscape did not include several of the most dominant species in our observational community composition plots (e.g. *Avena fatua*, *A. barbata*, *Bromus diandrus*, *Microseris douglasii*, and *Lolium perenne* each achieved >50% cover in at least one 1×1 plot across the 16 sites, but were not part of the experiment). Moreover, the functional traits of some of the most dominant species were beyond the range of functional traits captured by the 17 species in our demography experiment (e.g. all five of the aforementioned dominant species had SRL values below those of the 17 species in the experiment, Fig. S2). As CWM trait values are intrinsically reflective of dominant species’ responses to the landscape, it is possible that including more species that captured a wider range of the trait variation found in our community would reveal trait-environment interactions that drive trait turnover patterns at the community level.

A second limitation of our study is that we were unable to account for the possibility that intra-specific trait variation (ITV) driven by local adaptation, phenotypic plasticity, or maternal effects – processes that are known to be important in similar serpentine systems (Rajakaruna and Bohm 1999; Baythavong 2011; Germain and Gilbert 2014) – mediate trait-environment relations at either the community or individual scale. However, our finding that trait-performance relationships do change across the environmental gradient generate predictions for future studies about how ITV may be structured on this landscape. For example, our result finding that the optimal value of SLA shifts from low to high with an increase in soil Ca:Mg (Fig. 3E) suggests that ITV may be structured such that individuals of the same species growing in soils with higher Ca:Mg build higher-SLA leaves than conspecific individuals on low-Ca:Mg soils. Understanding how the spatial structure of ITV differs between species may be critical for predicting variation between species in their demographic responses to environmental gradients (Swenson et al. 2020).

## Conclusion

Understanding and forecasting how species and communities respond to environmental variation is a fundamental challenge in ecology. Predicting variation in species-level demographic processes based on patterns in trait turnover across whole communities is a promising approach, but most methods to do so have relied on the assumption that variation in community-weighted mean (CWM) traits reflect shifts in trait optima over landscapes. Our study found consistent evidence that variation in CWM traits across environmental gradients reflect the effects of changing trait-performance relationships, but they our results caution against inferring likely demographic responses of plants to environments on the basis of CWM traits alone. Future efforts that link plant traits to variation in population growth across variable environments rates will help build towards more predictive trait-based models of plant community dynamics.

## Acknowledgements

We acknowledge the Chumash peoples as the traditional land caretakers of the ecosystem studied in this project, and the Gabrielino/Tongva peoples as the traditional land caretakers of Tovaangar (the Los Angeles basin and So. Channel Islands), where UCLA is located. We are grateful to Jonathan Levine and Oscar Godoy for help in developing the experimental plot network used for this analyses. We thank Clare Camilleri, Anmol Dhaliwal, Aoefe Galvi, Renato Guidon, Jonathan Levine, Mirjam von Rutte, Mary Van Dyke, and Xinyi Yan for help in data collection, and Kate McCurdy and other staff at Sedgwick Reserve for help in the field. We thank Madeline Cowen, Kenji Hayashi, Mary Van Dyke, Marcel Vaz, and the DataPhiles group at the University of Missouri for comments on the analysis and manuscript. This work was performed in part at the University of California Natural Reserve System Sedgwick Reserve (DOI: 10.21973/N3C08R). This work was funded by the National Science Foundation DEB-1644641.

## Supplemental figures

**Figure S1:**
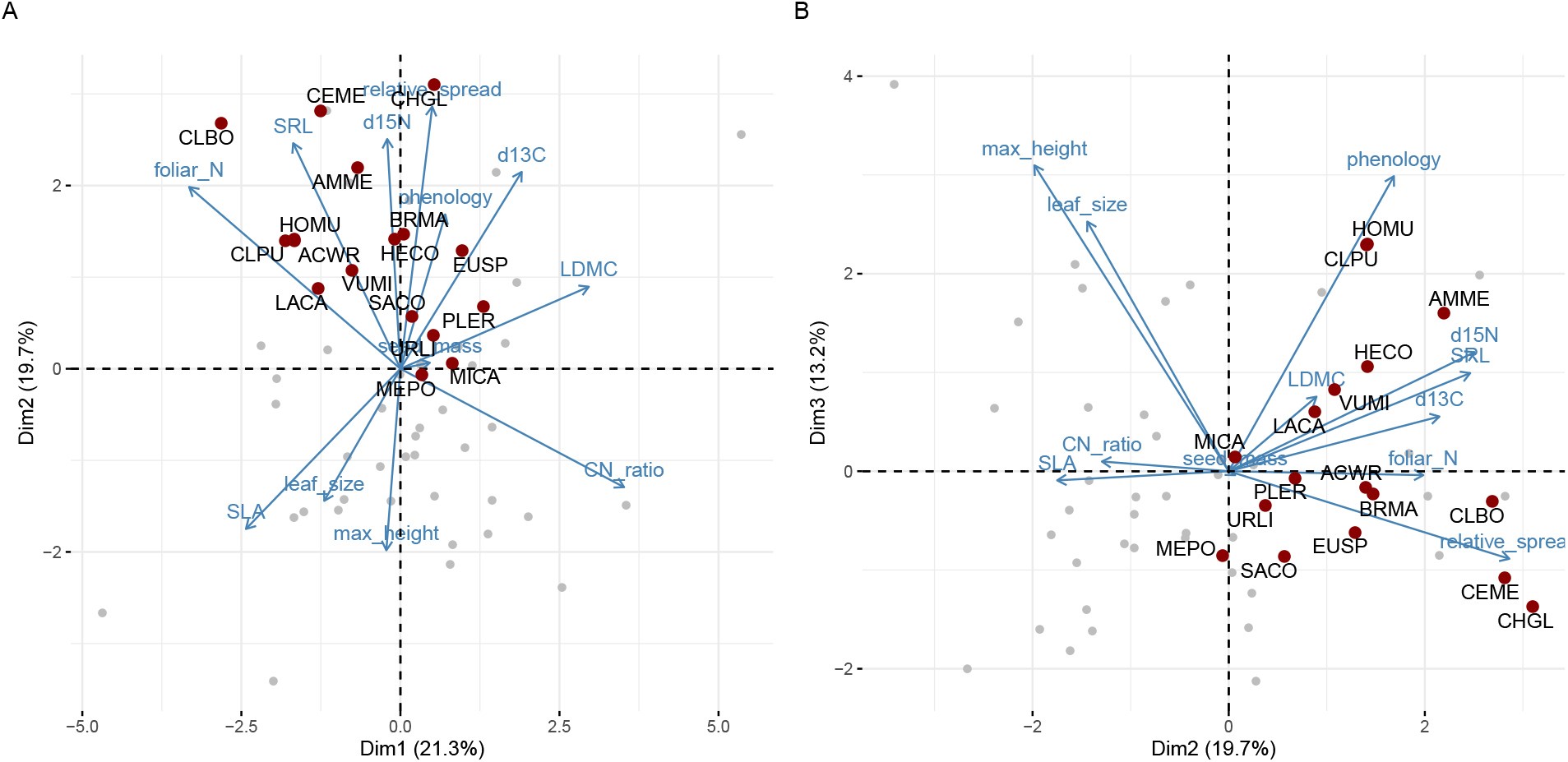
Biplots of axes 1/2 (Panel A) and axes 2/3 (Panel B) from a PCA of the functional traits measured for this study. Light grey points indicate the position of the species found across the community (N = 55), and red points indicate the position of each of the focal species of the demography experiment (N = 17)

**Figure S2:**
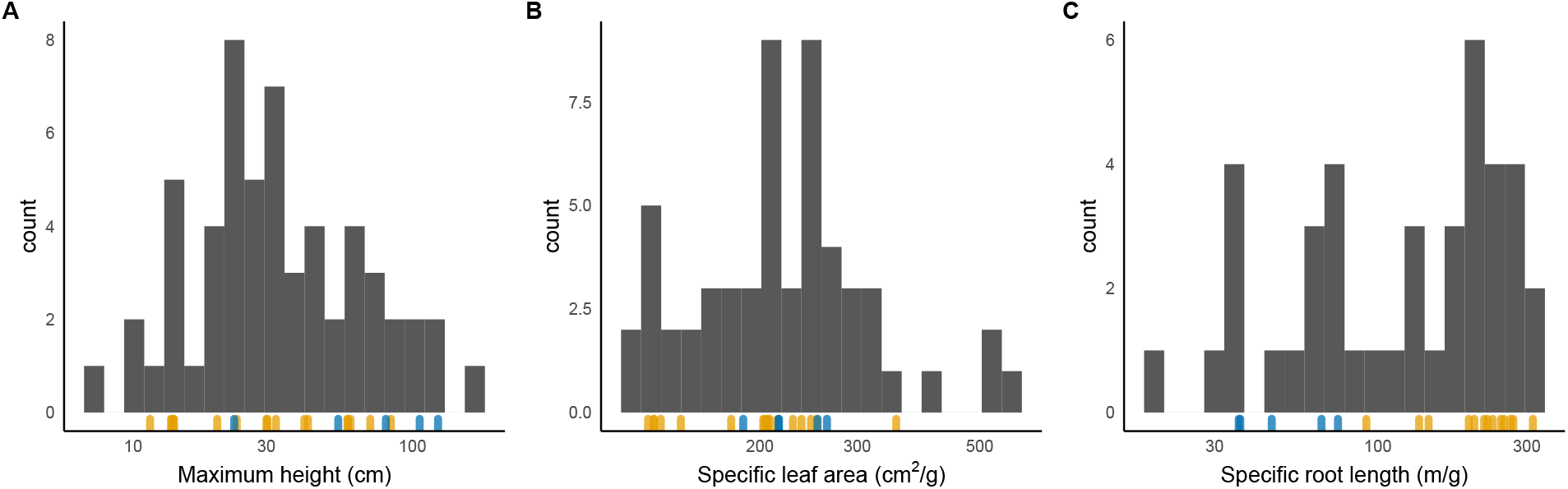
Histograms of the three focal for all species encountered in the Serpentine grassland at Sedgwick Reserve. Each blue tick at the bottom of the histograms indicates the trait value for one of the focal species in the demography experiment, and the orange ticks indicate trait values of species that were dominant in the community (relative cover > 50 in at least one site) but absent in our experiment. Note the log-transformed X-axis in each panel.

**Figure S3:**
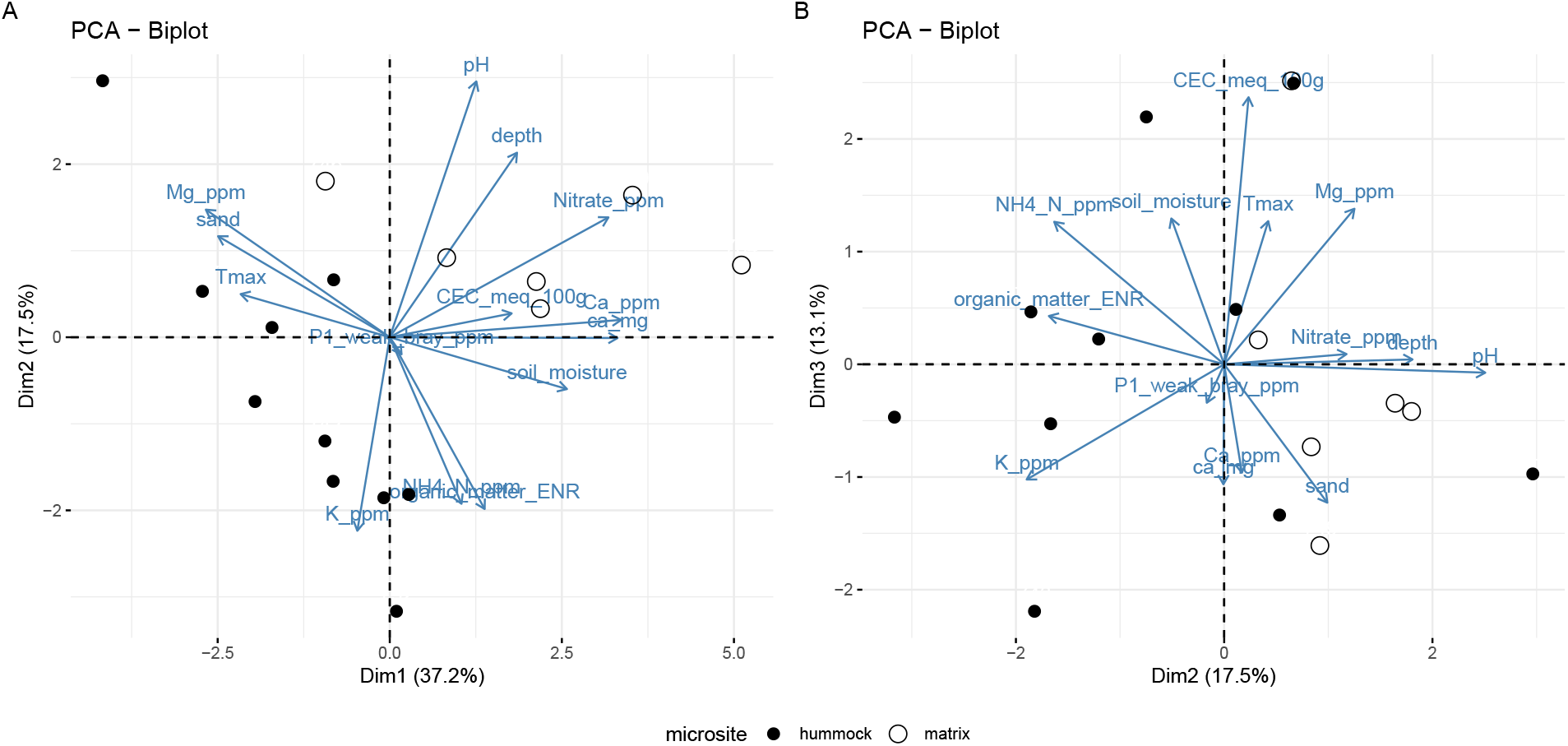
Biplots of axes 1/2 (Panel A) and axes 2/3 (Panel B) from a PCA of the environmental gradients measured for this study.

**Figure S4:**
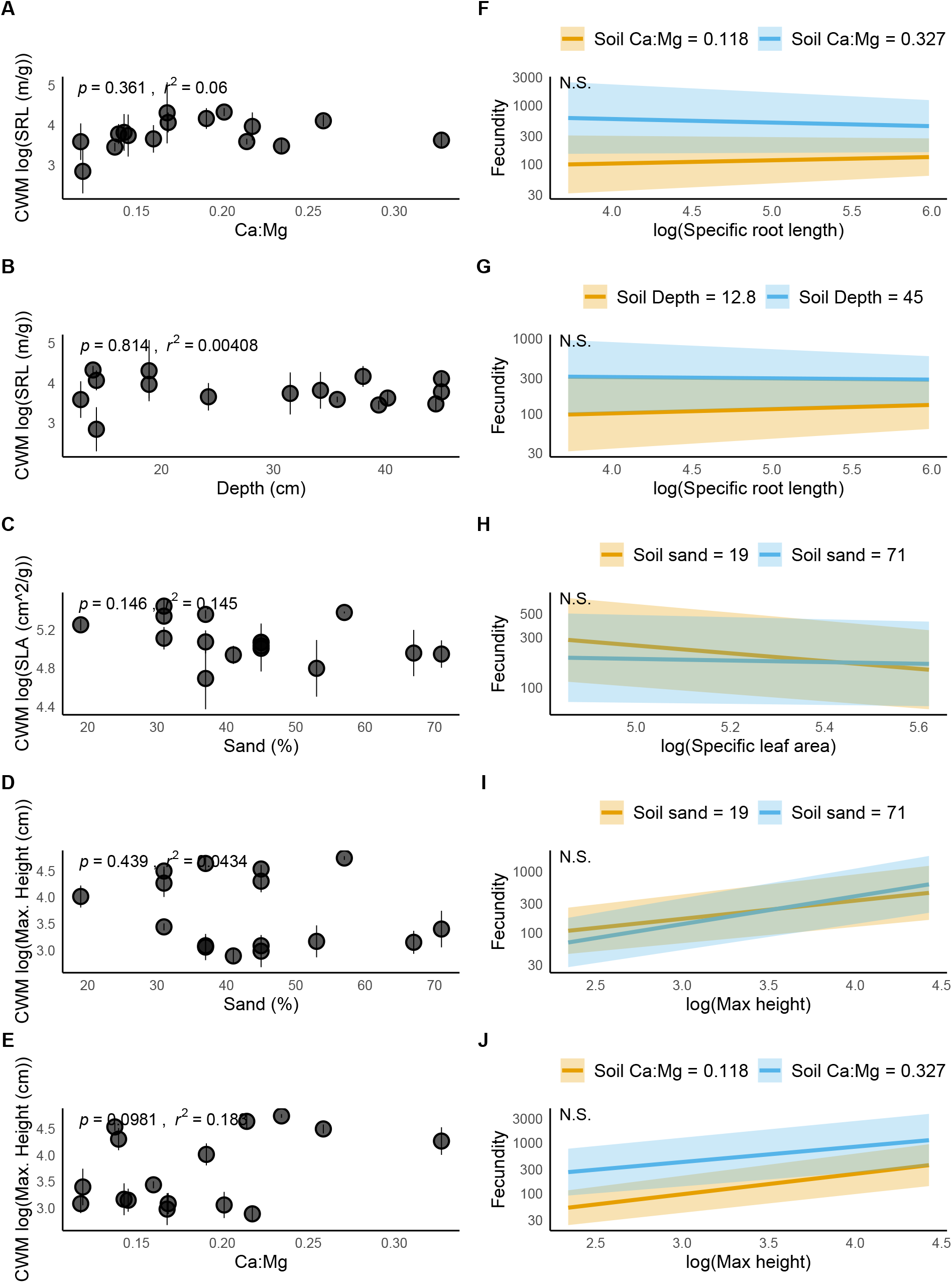
Panels A-E are biplots of CWM trait values and environmental variables for the five pairwise comparisons that were non-significant. Panels F-J show the corresponding trait-environment interactions in the model of seed output as a function of trait and23environmental predictors.

**Figure S5:**
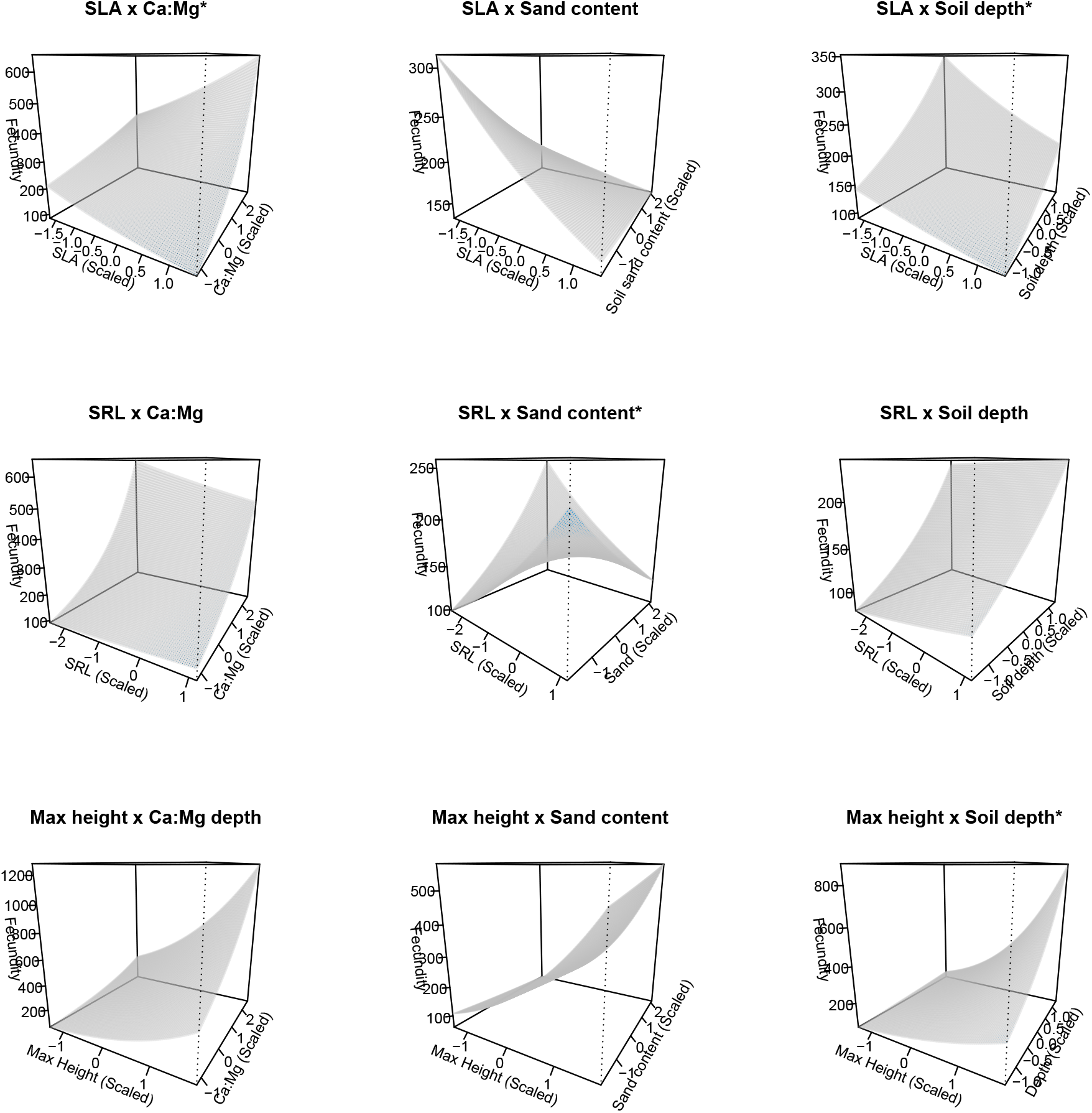
3D interaction surfaces for all nine trait-environment interactions in our GLMM of seed production as a function of trait and environment predictors. Plots labeled with an asterisk indicate significant interaction terms.

**Table S1:** List of traits measured for this study

| Organ | Trait | Units |
| --- | --- | --- |
| Whole plant | Max. height | cm |
|  | Canopy shape index | dimensionless |
|  | Carbon isotope composition (dC13) | dC13 |
|  | Phenology | day of year |
|  | Seed mass | mg |
| Leaf | Leaf size | cm<sup>2</sup> |
|  | Specific leaf area | g/cm<sup>2</sup> |
|  | Leaf dry matter content | mg/g |
|  | C:N ratio | dimensionless |
|  | Leaf N concentration | mg/g |
| Root | Specific root length | m/g |

**Table S2:**
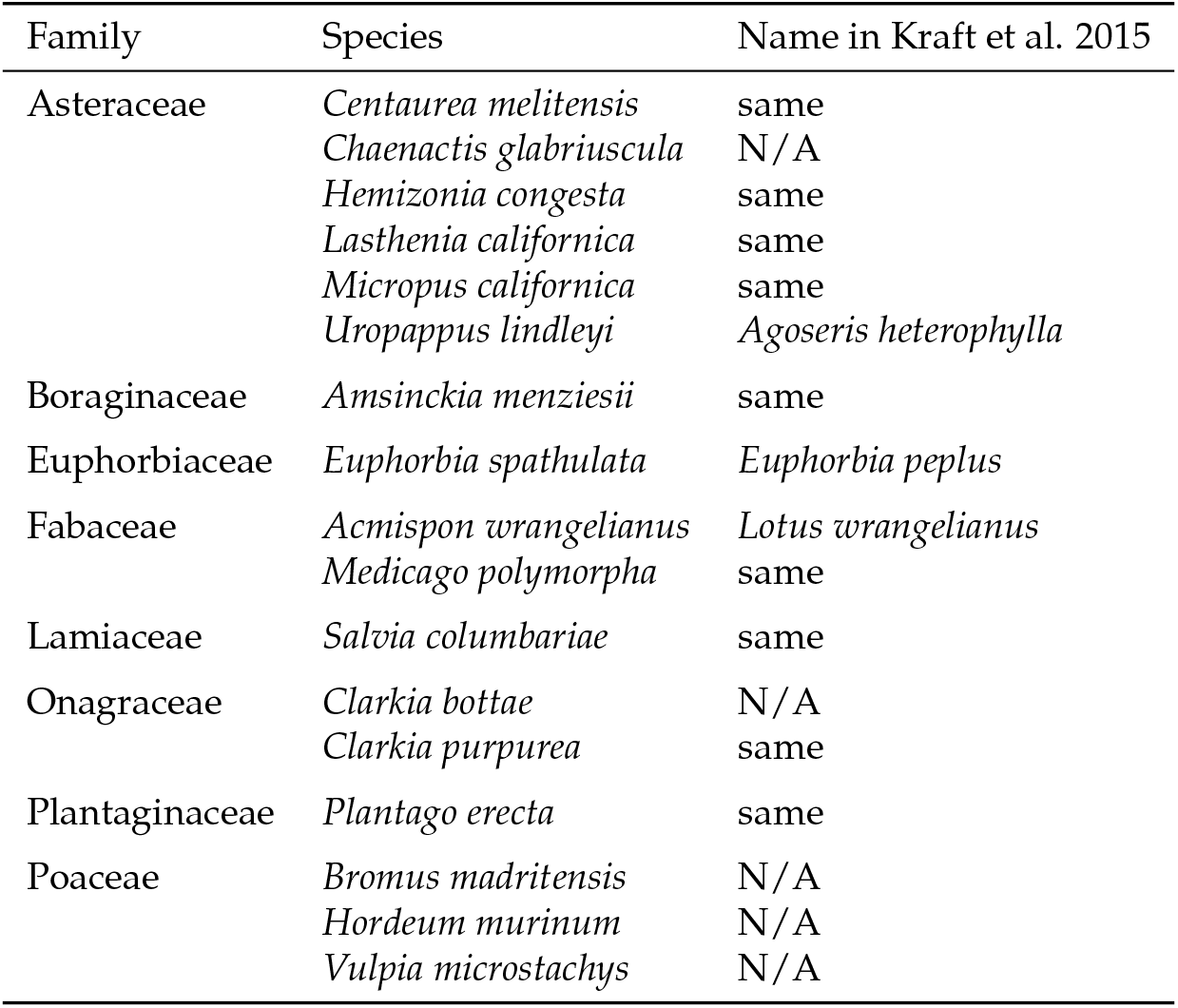
Species used in the demography experiment.

**Table S3:** Model output for quadratic relationships between CWM traits and environmental variables

| trait | environment | term | estimate | std.error | statistic | p.value |
| --- | --- | --- | --- | --- | --- | --- |
| SLA | Ca:Mg | Intercept | 4.1710630 | 0.3810 | 10.9473 | 0.0000 |
| SLA |  | Linear | 6.7997132 | 3.7945 | 1.7920 | 0.0964 |
| SLA |  | Quadratic | -8.8295749 | 8.8486 | -0.9978 | 0.3366 |
| SLA | Depth | Intercept | 4.8685302 | 0.3669 | 13.2676 | 0.0000 |
| SLA |  | Linear | 0.0039068 | 0.0296 | 0.1322 | 0.8968 |
| SLA |  | Quadratic | 0.0001119 | 0.0005 | 0.2187 | 0.8302 |
| SLA | Sand | Intercept | 5.6767840 | 0.5039 | 11.2652 | 0.0000 |
| SLA |  | Linear | -0.0211202 | 0.0225 | -0.9375 | 0.3656 |
| SLA |  | Quadratic | 0.0001628 | 0.0002 | 0.6820 | 0.5072 |
| SRL | Ca:Mg | Intercept | 1.3091538 | 0.9470 | 1.3824 | 0.1901 |
| SRL |  | Linear | 24.1034774 | 9.4310 | 2.5558 | 0.0239 |
| SRL |  | Quadratic | -53.0592298 | 21.9926 | -2.4126 | 0.0313 |
| SRL | Depth | Intercept | 3.7116730 | 0.8014 | 4.6316 | 0.0005 |
| SRL |  | Linear | 0.0093611 | 0.0645 | 0.1450 | 0.8869 |
| SRL |  | Quadratic | -0.0001975 | 0.0011 | -0.1768 | 0.8624 |
| SRL | Sand | Intercept | 3.6723606 | 0.7948 | 4.6207 | 0.0005 |
| SRL |  | Linear | 0.0207823 | 0.0355 | 0.5849 | 0.5686 |
| SRL |  | Quadratic | -0.0003831 | 0.0004 | -1.0175 | 0.3275 |
| Max. Height | Ca:Mg | Intercept | 2.9756663 | 1.9356 | 1.5373 | 0.1482 |
| Max. Height |  | Linear | 2.8657993 | 19.2769 | 0.1487 | 0.8841 |
| Max. Height |  | Quadratic | 5.5585773 | 44.9527 | 0.1237 | 0.9035 |
| Max. Height | Depth | Intercept | 3.4516879 | 0.7104 | 4.8586 | 0.0003 |
| Max. Height |  | Linear | -0.0486766 | 0.0572 | -0.8508 | 0.4103 |
| Max. Height |  | Quadratic | 0.0016727 | 0.0010 | 1.6891 | 0.1150 |
| Max. Height | Sand | Intercept | 4.4618081 | 1.7386 | 2.5663 | 0.0235 |
| Max. Height |  | Linear | -0.0246015 | 0.0777 | -0.3165 | 0.7566 |
| Max. Height |  | Quadratic | 0.0001513 | 0.0008 | 0.1837 | 0.8571 |

## Appendices

