## Appendices S1 and S2 for "Functional traits predict species responses to environmental variation in a California grassland annual plant community"

Last rendered: 17 Nov 2020

*Preprint for bioRxiv*

**Authors**: Gaurav S. Kandlikar[1,2], Andew R. Kleinhesselink[1], Nathan J.B. Kraft[1]

[1] Dept. of Ecology and Evolutionary Biology, University of California, Los Angeles, Los Angeles, CA.

[2] Present address: Division of Biological Sciences, University of Missouri, Columbia, MO.

**Author for correspondence**:

Gaurav Kandlikar:

**Coauthor contact information**:

Andrew R. Kleinhesselink:

Nathan J.B. Kraft:

### Appendices

#### Appendix S1: Converting field measurements of reproductive output to estimates of seed count per individual

For this study we quantified the seed output of 1632 plants in the field. When possible we directly counted the number of seeds on each focal individual when plants were at their maximum reproductive output, but for time constraits had to measure proxies of reproductive output (e.g. diamater of flower head) for several species. In this Appendix we describe how we converted these proxy measurements of reproductive output into estimates of seed counts for each of the species in our experiment.

For AMME, BRMA, CEME, EUPE, HECO, HOMU, MICA, PLER, and VUMI we could simply count the number of seeds on each focal plant in the field. For AGHE, LOWR, and MEPO, we counted the number of seed heads or fruits per individual in the field. We separately counted the number of seeds in 40 mature seed heads or fruits, and multiplied the fruit or seed head count by the mean count of seeds per fruit to estimate the number of seeds made by each plant in the field (Fig S1.1). For CHGL, LACA, SACO, we measured the diamater of each flower head in the field. We separately built a regression between the area of seed heads (calculated assuming the seed head was circular) and number of seeds (Fig S1.2), and used the regression equation to estimate the number of seeds produced by each species in the field. For CLBO and CLPU, we measured the length of each fruit in the field, and again used a regression equation between fruit length and number of seeds per fruit to calculate the seed production of focal individuals (Fig S1.2). We rounded each calculated seed count down to the nearest integer. When a focal individual was too large to count every seed, fruit or flower head, we counted the reproductive units on a quarter or half of the plant, and multiplied as appropriate for a whole-plant estimate.

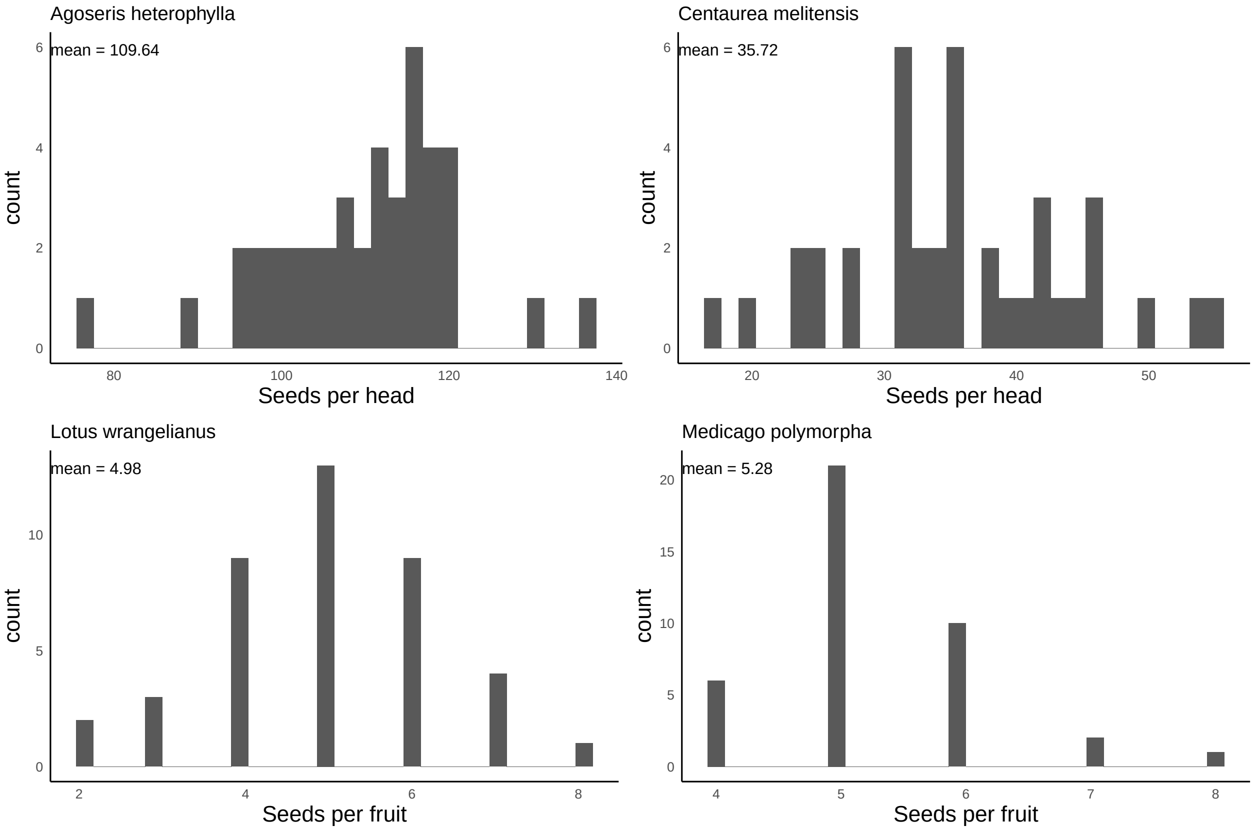


Figure S1.1: Histogram of the number of seeds per flower head (or fruit) for Agoseris heterophylla, Centaurea melitensis, Lotus wrangelianus, and Medicago polymorpha

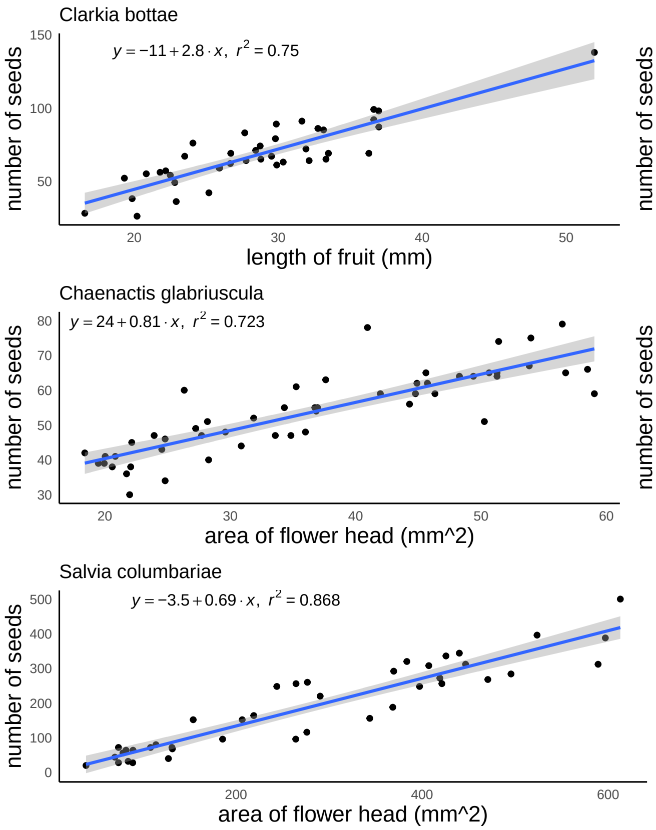


Figure S1.2: Regressions of seed count as a function of flower head or fruit size for Clarkia bottae, Clarkia purpurea, Chaenatcis glabriuscula, Lasthenia californica, and Salvia columbariae

### Appendix S2: R Code to recreate all analyses in the manuscript

#### Setup

```
# All packages from CRAN:
library(bbmle)
library(broom)
library(broom.mixed)
library(DHARMa)
library(effects)
library(factoextra)
library(FactoMineR)
library(glmmTMB)
library(latex2exp)
library(patchwork)
library(tidyverse)
library(visreg)

# Import sedgwickcover libraries
# devtools::install_github("akleinhesselink/sedgwickcover")
library(sedgwickcover)
theme_plots <- function() {
  theme_minimal() +
    theme(panel.border = element_blank(),
                   panel.grid.major = element_blank(),
                   panel.grid.minor = element_blank(),
                   axis.line = element_line(colour = "black"),
                   plot.tag = element_text(face = "bold"),
                   axis.title = element_text(size = 12),
                   plot.caption = element_text(size = 12)
    )
}
theme_set(theme_plots())
```

#### Community-weighted trait means

Import trait and cover data

```r
strts_all <- read_csv("data/trait/all_trait_data.csv") %>% select(-X1)
```

```
## Warning: Missing column names filled in: 'X1' [1]
```

```
## Parsed with column specification:
## cols(
##   .default = col_double(),
##   USDA_symbol = col_character(),
##   notes = col_character(),
##   seed_mass_data_source = col_character(),
##   max_height_data_source = col_character(),
##   dataset = col_character()
## )
```

```
## See spec(...) for full column specifications.
```

```r
strts_all %>%
  group_by(USDA_symbol) %>%
  summarize(n = n()) %>%
  arrange(-n)
```

```
## `summarise()` ungrouping output (override with `.groups` argument)
```

```
## # A tibble: 57 x 2
##    USDA_symbol     n
##    <chr>       <int>
##  1 ANAR            2
##  2 BRMA3           2
##  3 CHGL            2
##  4 FEMI2           2
##  5 HOMU            2
##  6 MICA            2
##  7 MIDO            2
##  8 SIGA            2
##  9 ACMO2           1
## 10 AGHE2           1
## # ... with 47 more rows
```

```r
# Remove some duplicate rows from the traits df
# (These species were measured in multiple years;
```

```r
# retaining only those measurements made in 2017, i.e.
# the year of the community cover sampling).
strts <- strts_all %>%
  group_by(USDA_symbol) %>%
  filter(!(USDA_symbol == "ANAR" & dataset == "TAPIOCA")) %>%
  filter(!(USDA_symbol == "BRMA3" & dataset == "2016")) %>%
  filter(!(USDA_symbol == "CHGL" & dataset == "2016")) %>%
  filter(!(USDA_symbol == "FEMI2" & dataset == "2016")) %>%
  filter(!(USDA_symbol == "HOMU" & dataset == "2016")) %>%
  filter(!(USDA_symbol == "MICA" & dataset == "2016")) %>%
  filter(!(USDA_symbol == "MIDO" & dataset == "TAPIOCA")) %>%
  filter(!(USDA_symbol == "SIGA" & dataset == "TAPIOCA")) %>%
  return




# Import the cover data
p17 <- sedgwickcover::plot_cover_2017

# Calculate the relative cover in each plot at each site
p17 <- p17 %>%
  filter(site < 756) %>%
  group_by(site, plot) %>%
  summarize(total_cover = sum(cover)) %>%
  left_join(p17, .) %>%
  mutate(relative_cover = cover/total_cover) %>%
  as_tibble()
```

```
## ‘summarise()‘ regrouping output by ’site’ (override with ‘.groups‘ argument)
## Joining, by = c("site", "plot")
```

```r
# Merge the cover and trait data frames
p17_trt <- left_join(p17, strts)
```

```
## Joining, by = "USDA_symbol"
```

```r
# species with missing data for at least one of the traits
missing_trts <-
```

```r
  p17_trt %>%
  group_by(USDA_symbol, site, plot) %>%
  filter(is.na(SLA) | is.na(SRL) | is.na(max_height)) %>%
  arrange(-relative_cover)

missing_trts_nz <- missing_trts %>% filter(relative_cover > 0)
mean(missing_trts_nz$relative_cover)
```

```
## [1] 0.04041392
```

```r
# std error function
sem <- function(x) (sd(x)/sqrt(n()))

# Summarize the trait data to begin calculating CWM mean
# (and sd of CWM trait) at each site
p17_trt_sum <-
  p17_trt %>%
  mutate(lsla = (log(SLA)),
         lsrl = (log(SRL)),
         lmaxht = (log(max_height))) %>%
  select(site:relative_cover, lsla, lsrl, lmaxht) %>%
  mutate(lsla_contrib = lsla*relative_cover,
         lsrl_contrib  = lsrl*relative_cover,
         lmaxht_contrib = lmaxht*relative_cover) %>%
  group_by(site, plot) %>%
  summarize_at(c("lsla_contrib", "lsrl_contrib", "lmaxht_contrib"),
               sum, na.rm = T) %>%
  rename(lsla_cwm = lsla_contrib,
         lsrl_cwm = lsrl_contrib,
         lmaxht_cwm = lmaxht_contrib)

# Summarize across the sub-plots at each site, to get the
# mean CWM, and SD+SEM at each site
p17_sitesum <-
  p17_trt_sum %>%
  group_by(site) %>%
  summarize_at(c("lsla_cwm", "lsrl_cwm", "lmaxht_cwm"),
```

```r
               lst(mean, sd, sem))


env <- read_csv("data/environmental/all_environmental_data.csv")
```

```
## Parsed with column specification:
## cols(
##   .default = col_double(),
##   type = col_character(),
##   microsite = col_character()
## )
## See spec(...) for full column specifications.
```

```r
env$ca_mg <- env$Ca_ppm/env$Mg_ppm
# merge in the environmental data with both plot-level and site-level CWM dfs
p17_sitesum_env <- env %>%
  rename(site = plot) %>%
  left_join(p17_sitesum, .)
```

```
## Joining, by = "site"
```

```r
# Remove the lower reserve sites, which got added again in the join
p17_sitesum_env <- p17_sitesum_env %>% filter(site < 756)
```

Now, compute CWM-environment relations:

```r
# Now, test for CWM trait shifts along the focal environmental axes ----
# Import functions that run LMs and make plots
source("analyses/functions/make_trait_env_lms.R")
source("analyses/functions/make_trait_env_plot.R")
lsla_sand_lm <- make_trait_env_lms(trait = "lsla_cwm_mean", env = "sand")
lsla_sand_plot <- make_trait_env_plot(trait = "lsla_cwm_mean", env = "sand",
                                       lsla_sand_lm, xl = "Sand (%)", yl = "CWM log(SLA (cm^2/g))")
lsla_camg_lm <- make_trait_env_lms(trait = "lsla_cwm_mean", env = "ca_mg")
lsla_camg_plot <- make_trait_env_plot(trait = "lsla_cwm_mean", env = "ca_mg",
                                       lsla_camg_lm, xl = "Ca:Mg", yl = "CWM log(SLA (cm^2/g))")
lsla_depth_lm <- make_trait_env_lms(trait = "lsla_cwm_mean", env = "depth")
lsla_depth_plot <- make_trait_env_plot(trait = "lsla_cwm_mean", env = "depth",
                                        lsla_depth_lm, xl = "Depth (cm)", yl = "CWM log(SLA (cm^2/g))")
```

```
lsrl_sand_lm <- make_trait_env_lms(trait = "lsrl_cwm_mean", env = "sand")
lsrl_sand_plot <- make_trait_env_plot(trait = "lsrl_cwm_mean", env = "sand",
                                       lsrl_sand_lm, xl = "Sand (%)", yl = "CWM log(SRL (m/g))")
lsrl_camg_lm <- make_trait_env_lms(trait = "lsrl_cwm_mean", env = "ca_mg")
lsrl_camg_plot <- make_trait_env_plot(trait = "lsrl_cwm_mean", env = "ca_mg",
                                       lsrl_camg_lm, xl = "Ca:Mg", yl = "CWM log(SRL (m/g))")
lsrl_depth_lm <- make_trait_env_lms(trait = "lsrl_cwm_mean", env = "depth")
lsrl_depth_plot <- make_trait_env_plot(trait = "lsrl_cwm_mean", env = "depth",
                                        lsrl_depth_lm, xl = "Depth (cm)", yl = "CWM log(SRL (m/g))")

lmaxht_sand_lm <- make_trait_env_lms(trait = "lmaxht_cwm_mean", env = "sand")
lmaxht_sand_plot <- make_trait_env_plot(trait = "lmaxht_cwm_mean", env = "sand",
                                         lmaxht_sand_lm, xl = "Sand (%)", yl = "CWM log(Max. Height (cm
lmaxht_camg_lm <- make_trait_env_lms(trait = "lmaxht_cwm_mean", env = "ca_mg")
lmaxht_camg_plot <- make_trait_env_plot(trait = "lmaxht_cwm_mean", env = "ca_mg",
                                         lmaxht_camg_lm, xl = "Ca:Mg", yl = "CWM log(Max. Height (cm))"
lmaxht_depth_lm <- make_trait_env_lms(trait = "lmaxht_cwm_mean", env = "depth")
lmaxht_depth_plot <- make_trait_env_plot(trait = "lmaxht_cwm_mean", env = "depth",
                                          lmaxht_depth_lm, xl = "Depth (cm)", yl = "CWM log(Max. Height

cwmtrts_plot <- wrap_plots(list(lsla_sand_plot,  lsla_camg_plot, lsla_depth_plot,
                                lsrl_sand_plot,  lsrl_camg_plot, lsrl_depth_plot,
                                lmaxht_sand_plot,lmaxht_camg_plot, lmaxht_depth_plot), nrow = 3)
cwmtrts_plot
```

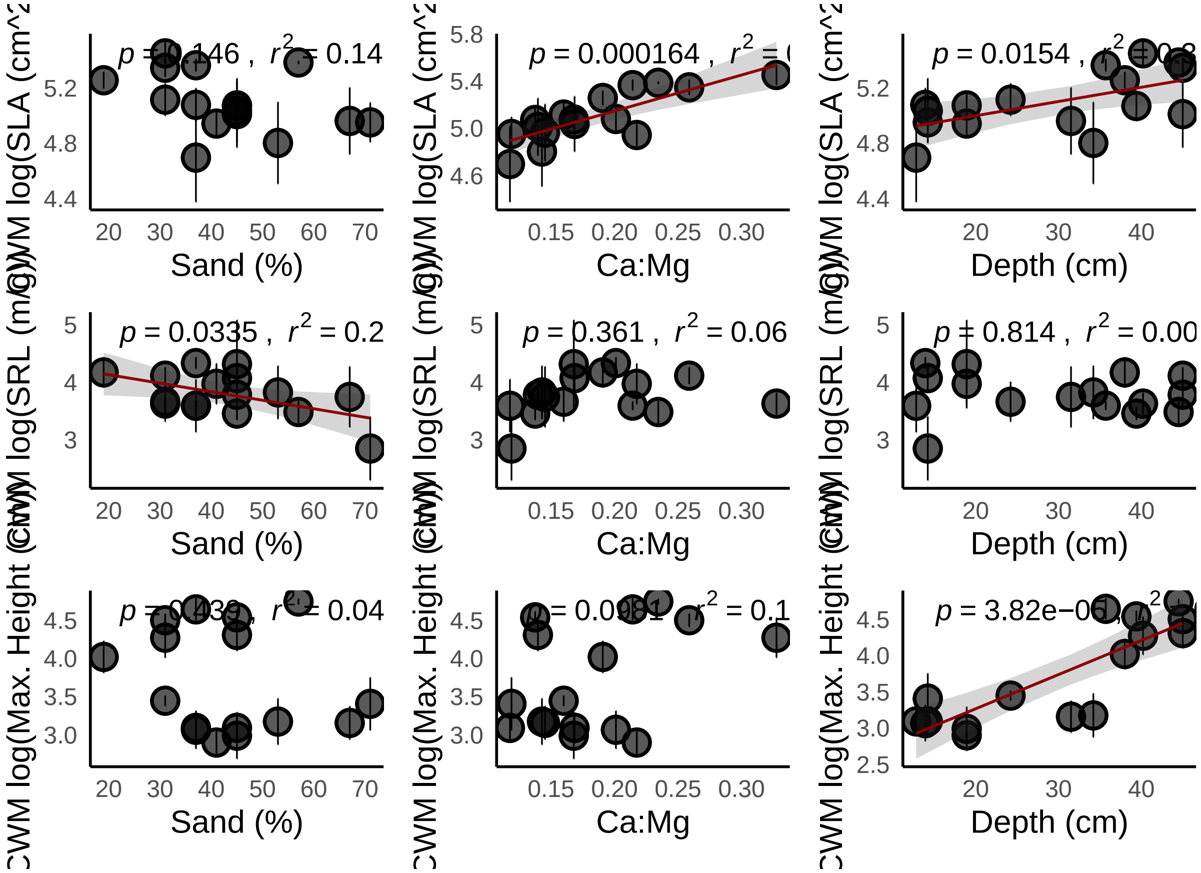


There are four significant CWM trait-environment relationships: SLA-Ca:Mg; SLA:Depth; SRL:Sand; Max. Height:Depth

#### Demography experiment

Import performance data, merge with trait data, and merge with environmental variables data.

```
## Now, we can test for trait-environment interactions in the demography data ----


# Clean/manage performance data ------
performance <- read_csv("data/performance/seed_production_processed.csv")
```

```
## Parsed with column specification:
## cols(
##   plot_num = col_double(),
##   plot_type = col_character(),
##   replicate = col_double(),
##   sp_code = col_character(),
##   num_seeds_produced = col_double()
```

```
89 ## )
```

```
performance <- performance  %>%
  # Include only the no-competition individuals
  filter(plot_type == "L") %>%
  # NA in this means the seed production was zero
  mutate(num_seeds_produced = ifelse(is.na(num_seeds_produced), 0,
                                     num_seeds_produced))
performance <- performance %>%
  # Make a new column for Hummock or matrix
  mutate(plot_env = ifelse(plot_num %in% c(742:745,747,748,750,751,752,755),
                     "hummock", "matrix")) %>%
  # remove lower-reserve plots
  filter(plot_num < 756) %>%
  return

# Make a new df to help match species names between trait and performance dfs
sp_code_usda_symbol <- data.frame(
  sp_code = unique(performance$sp_code),
  USDA_symbol = c("URLI5", "AMME", "BRMA3", "CEME2",
                 "CLBO", "CLPU2", "CHGL", "EUSP",
                 "HECO7", "HOMU", "LACA7", "LOWR2",
                 "MEPO3", "MICA", "PLER3", "SACO6", "FEMI2")
)

performance <- left_join(performance, sp_code_usda_symbol)
```

```
90 ## Joining, by = "sp_code"
```

```
# Add in the environmental data
perf_env <- left_join(performance,
                      env %>%
                        # keep only the focal environmental variables, for now
                        select(plot_num = plot, unscaled_ca_mg = ca_mg,
                               unscaled_sand = sand, unscaled_depth = depth))
```

```
91 ## Joining, by = "plot_num"
```

```r
perf_env
```

```
## # A tibble: 1,399 x 10
##    plot_num plot_type replicate sp_code num_seeds_produ~ plot_env USDA_symbol
##       <dbl> <chr>         <dbl> <chr>              <dbl> <chr>    <fct>
##  1      740 L                 1 AGHE                 768 matrix   URLI5
##  2      740 L                 2 AGHE                1206 matrix   URLI5
##  3      740 L                 3 AGHE                1425 matrix   URLI5
##  4      740 L                 4 AGHE                1864 matrix   URLI5
##  5      740 L                 5 AGHE                1425 matrix   URLI5
##  6      740 L                 6 AGHE                 987 matrix   URLI5
##  7      741 L                 1 AGHE                1206 matrix   URLI5
##  8      741 L                 2 AGHE                 987 matrix   URLI5
##  9      741 L                 3 AGHE                 548 matrix   URLI5
## 10      741 L                 4 AGHE                1206 matrix   URLI5
## # ... with 1,389 more rows, and 3 more variables: unscaled_ca_mg <dbl>,
## #   unscaled_sand <dbl>, unscaled_depth <dbl>
```

```r
# Add in the trait data

# Include only the trait data collected in 2016 or for TAPIOCA
trait_df16 <-
  strts_all %>%
  filter(dataset %in% c("TAPIOCA", "2016")) %>%
  select(USDA_symbol, SLA, SRL, max_height)

perf_env_trt <- left_join(perf_env, trait_df16, by = "USDA_symbol")

perf_env_trt$unscaled_log_sla <- log(perf_env_trt$SLA)
perf_env_trt$unscaled_log_srl <- log(perf_env_trt$SRL)
perf_env_trt$unscaled_log_maxht <- log(perf_env_trt$max_height)

# perf_env_trt <- left_join(perf_env, traits)
perf_env_trt$plot_num <- as.factor(perf_env_trt$plot_num)
perf_env_trt$num_seeds_produced <- as.integer(perf_env_trt$num_seeds_produced)
```

Now, run the trait-environment GLMMs

```r
# Trait-Env model based on CWM trait ----
# SLA-Ca:Mg; SLA:Depth; SRL:Sand; Max. Height:Depth

# scale the relavent columns
perf_env_trt <- perf_env_trt %>% mutate(
  ca_mg = scale(unscaled_ca_mg),
  depth = scale(unscaled_depth),
  sand = scale(unscaled_sand),
  log_sla = scale(unscaled_log_sla),
  log_srl = scale(unscaled_log_srl),
  log_maxht = scale(unscaled_log_maxht))

trait_model_full <- glmmTMB(num_seeds_produced ~
                              ca_mg* log_sla +
                              depth*log_sla +
                              sand*log_sla +
                              ca_mg*log_srl +
                              depth*log_srl +
                              sand*log_srl +
                              ca_mg*log_maxht +
                              sand*log_maxht +
                              depth*log_maxht +
                              (1|sp_code) + (1|plot_num),
                            ziformula = ~1,
                            data = perf_env_trt,
                            family = nbinom2(link = "log"))
summary(trait_model_full)
```

```
##  Family: nbinom2  ( log )
## Formula:
## num_seeds_produced ~ ca_mg * log_sla + depth * log_sla + sand *
##     log_sla + ca_mg * log_srl + depth * log_srl + sand * log_srl +
##     ca_mg * log_maxht + sand * log_maxht + depth * log_maxht +
##     (1 | sp_code) + (1 | plot_num)
## Zero inflation:                      ~1
## Data: perf_env_trt
```

```
##
##      AIC      BIC   logLik deviance df.resid
##  13257.2  13362.1  -6608.6  13217.2     1379
##
## Random effects:
##
## Conditional model:
##  Groups   Name        Variance Std.Dev.
##  sp_code  (Intercept) 0.5397   0.7346
##  plot_num (Intercept) 0.2423   0.4922
## Number of obs: 1399, groups:  sp_code, 17; plot_num, 16
##
## Overdispersion parameter for nbinom2 family (): 1.52
##
## Conditional model:
##                 Estimate Std. Error z value Pr(>|z|)
## (Intercept)      5.25146    0.22069  23.796  < 2e-16 ***
## ca_mg            0.36700    0.15281   2.402 0.016317 *
## log_sla         -0.13120    0.18216  -0.720 0.471371
## depth            0.33455    0.13897   2.407 0.016066 *
## sand            -0.02849    0.14056  -0.203 0.839352
## log_srl          0.02658    0.19626   0.135 0.892269
## log_maxht        0.55300    0.19739   2.802 0.005085 **
## ca_mg:log_sla    0.12470    0.03318   3.758 0.000171 ***
## log_sla:depth   -0.01903    0.02896  -0.657 0.511156
## log_sla:sand     0.04172    0.03079   1.355 0.175500
## ca_mg:log_srl   -0.04375    0.04193  -1.043 0.296795
## depth:log_srl   -0.03937    0.03556  -1.107 0.268229
## sand:log_srl    -0.10862    0.03462  -3.137 0.001707 **
## ca_mg:log_maxht -0.03911    0.04070  -0.961 0.336533
## sand:log_maxht   0.05951    0.03857   1.543 0.122862
## depth:log_maxht  0.09096    0.04050   2.246 0.024719 *
## ---
## Signif. codes:  0 '***' 0.001 '**' 0.01 '*' 0.05 '.' 0.1 ' ' 1
##
## Zero-inflation model:
```

```
##             Estimate Std. Error z value Pr(>|z|)
## (Intercept) -0.53232    0.05571  -9.556   <2e-16 ***
## ---
## Signif. codes:  0 '***' 0.001 '**' 0.01 '*' 0.05 '.' 0.1 ' ' 1
```

```
trait_model_null <- glmmTMB(num_seeds_produced ~
                              1 +
                              (1|sp_code) + (1|plot_num),
                            ziformula = ~1,
                            data = perf_env_trt,
                            family = nbinom2(link = "log"))



trait_model_additive <- glmmTMB(num_seeds_produced ~
                                  ca_mg + sand + depth +
                                  log_sla + log_srl + log_maxht +
                                  (1|sp_code) + (1|plot_num),
                                ziformula = ~1,
                                data = perf_env_trt,
                                family = nbinom2(link = "log"))
```

Check model performance:

```
# Verify whether interaction terms are supported
# relative to a model with only additive terms
# and a null model
bbmle::AICctab(trait_model_null, trait_model_full, trait_model_additive)
```

```
##                      dAICc df
## trait_model_full      0.0  20
## trait_model_additive 12.3  11
## trait_model_null     20.8  5
```

```
# Check VIF in the conditional component of the full model
performance::check_collinearity(trait_model_full, component = "conditional")
```

```
## # Check for Multicollinearity
##
## Low Correlation
##
```

```
##          Parameter  VIF Increased SE
##              ca_mg 1.46         1.21
##            log_sla 1.01         1.00
##              depth 1.20         1.10
##               sand 1.25         1.12
##            log_srl 1.18         1.08
##          log_maxht 1.18         1.09
##      ca_mg:log_sla 1.27         1.13
##      log_sla:depth 1.13         1.06
##       log_sla:sand 1.22         1.10
##      ca_mg:log_srl 1.55         1.25
##      depth:log_srl 1.48         1.22
##       sand:log_srl 1.50         1.23
##    ca_mg:log_maxht 1.73         1.31
##     sand:log_maxht 1.52         1.23
##    depth:log_maxht 1.61         1.27
```

```r
# Check other model diagnostics using DHARMa
resids <- DHARMa::simulateResiduals(trait_model_full)
plot(resids)
```

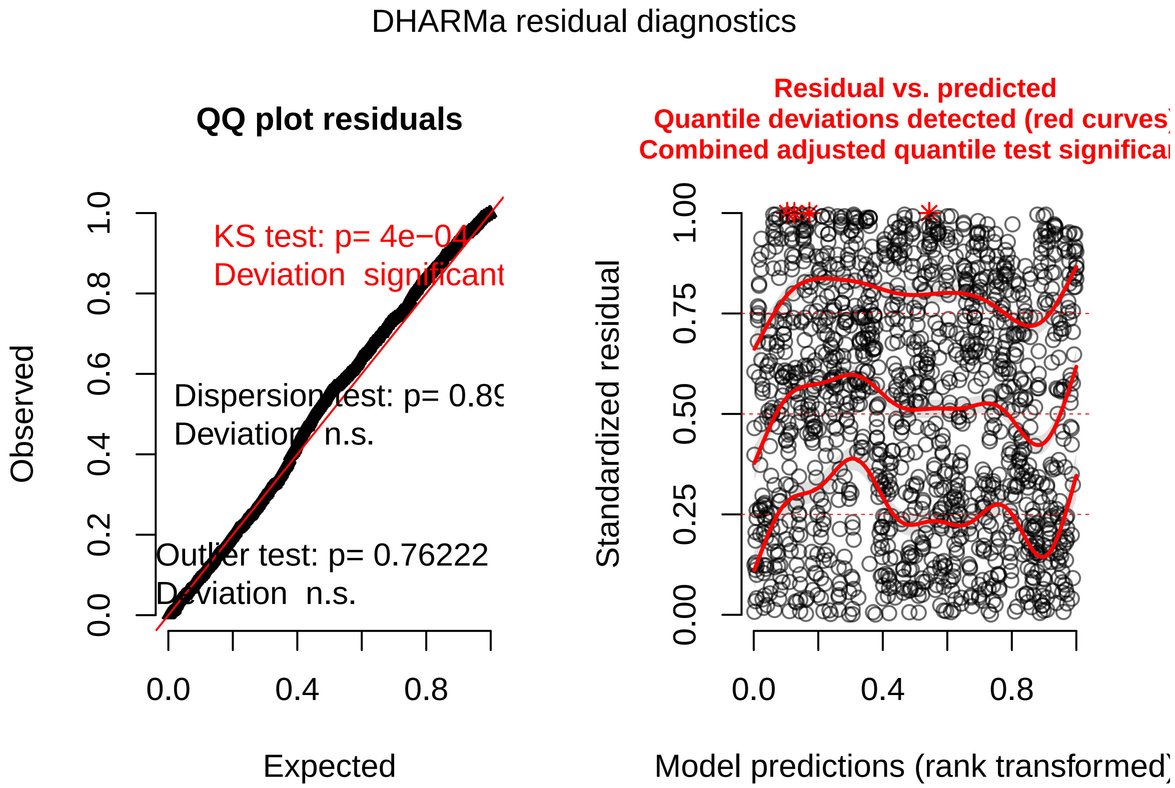

```r
# Looks some issue with uniformity - though the QQ looks fairly
# linear, and Florian Hartig's notes (e.g. https://github.com/florianhartig/DHARMa/issues/181)
# suggest that significance of Kolmogorov-Smirnov test need not be an
# issue with large numbers of samples.

## Compute the conditional and marginal R2
performance::r2_nakagawa(trait_model_full)
```

```
## # R2 for Mixed Models
##
##   Conditional R2: 0.372
##      Marginal R2: 0.178
```

Now, make plots

```r
## plot model coefficients ##

model_terms_full <-
  data.frame(confint(trait_model_full)) %>%
  rownames_to_column() %>%
  rename(term = rowname, lower_bound = X2.5..,
         upper_bound = X97.5.., estimate = Estimate) %>%
  filter(str_detect(string = term, pattern  = "cond.")) %>%
  filter(!str_detect(string = term, pattern  = "Intercept")) %>%
  mutate(term = str_replace(term, "cond.", ""),
         order = c(1,4,2,3,5:13,15,14)) %>%
  arrange(order) %>%
  mutate(term_clean = c("Soil Ca:Mg", "Soil depth", "Sand content",
                        "Specific leaf area (SLA)", "Specific root length (SRL)", "Max height",
                        "Ca:Mg x SLA", "Depth x SLA", "Sand content x SLA",
                        "Ca:Mg x SRL", "Depth x SRL", "Sand content x SRL",
                        "Ca:Mg x Max height", "Depth x Max height", "Sand content x Max height")) %>%
  arrange(-order) %>%
  mutate(term_clean = fct_reorder(term_clean, -order),
         fill_color = ifelse(upper_bound > 0 & lower_bound < 0, "NS", "S"))

(plot_full_model_coefficients <-
    ggplot(model_terms_full) +
```

```r
  geom_errorbar(aes(x = term_clean, ymin = lower_bound,
                    ymax = upper_bound),  color = alpha("grey", 0)) +
  geom_vline(xintercept = c(2,4,8,9), size = 10, color = "grey90") +
  geom_errorbar(aes(x = term_clean, ymin = lower_bound,
                    ymax = upper_bound), width = 0) +
  geom_point(aes(x = term_clean, y = estimate, fill = fill_color),
             size = 3, shape = 21, stroke = 1.5) +
  scale_fill_manual(values = c("white", "grey60")) +
  geom_hline(yintercept = 0, linetype = "dashed") +
  xlab("") +
  coord_flip() +
  ecoevoapps::theme_apps() +
  theme(axis.text = element_text(color = "black", size = 11),
        axis.title = element_text(size = 12),
        legend.position = "NA"))
```

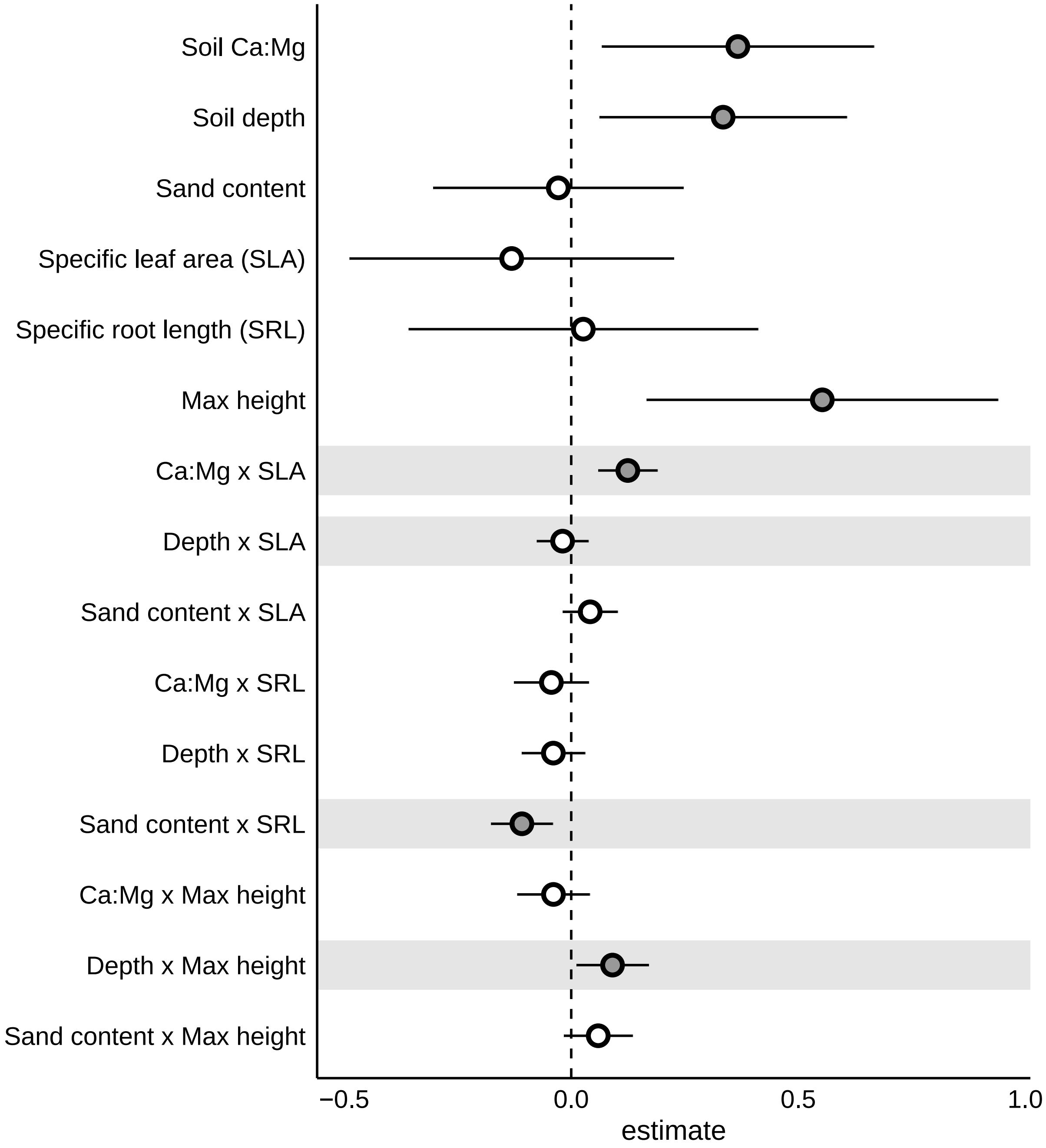


```
ggsave("manuscript/figures/model-coefficients.pdf", plot_full_model_coefficients,
       height = 6, width = 5)
```

```
## Interaction plots ##


colpal <- c( "#E69F00", "#56B4E9")


get_pval <- function(model_out = trait_model_full, mod_term) {
```

```r
  pval <- model_out %>% broom.mixed::tidy() %>%
    filter(term == mod_term) %>%
    select(p.value) %>% unlist

  pval <- ifelse(pval < .05,
                 paste0("\t p = ", round(pval, 4)),
                 "\t N.S.")
  pval
}

# make variables to help un-scale the axes
sla_sd <- sd(perf_env_trt$unscaled_log_sla)
sla_mean <- mean(perf_env_trt$unscaled_log_sla)
srl_sd <- sd(perf_env_trt$unscaled_log_srl)
srl_mean <- mean(perf_env_trt$unscaled_log_srl)
maxht_sd <- sd(perf_env_trt$unscaled_log_maxht)
maxht_mean <- mean(perf_env_trt$unscaled_log_maxht)

ca_mg_min <- round(min(perf_env_trt$unscaled_ca_mg), 3)
ca_mg_max <- round(max(perf_env_trt$unscaled_ca_mg), 3)
sand_min  <- round(min(perf_env_trt$unscaled_sand), 3)
sand_max  <- round(max(perf_env_trt$unscaled_sand), 3)
depth_min <- round(min(perf_env_trt$unscaled_depth), 3)
depth_max <- round(max(perf_env_trt$unscaled_depth), 3)

eff_sla_camg_df <- effects::effect("ca_mg:log_sla", trait_model_full,
                                xlevels = list(ca_mg = c(min(perf_env_trt$ca_mg),
                                                         max(perf_env_trt$ca_mg)),
                                               log_sla = c(min(perf_env_trt$log_sla),
                                                          max(perf_env_trt$log_sla)))) %>%
  data.frame
# The warning message can be ignored according to Ben Bolker in the GLMMTMB docs:
# https://github.com/glmmTMB/glmmTMB/blob/master/glmmTMB/inst/doc/model_evaluation.pdf
plot_sla_camg <-
    ggplot(eff_sla_camg_df, aes(x = (log_sla*sla_sd)+sla_mean, y = fit, color=factor(ca_mg)))+
    geom_line(size=1.25)+
```

```
    geom_ribbon(aes(ymin=lower,ymax=upper, fill=factor(ca_mg)),
                alpha=0.3, colour= NA) +
    annotate("text", x = -Inf, y = Inf, hjust = 0, vjust = 1,
             label = get_pval(mod_term = "ca_mg:log_sla")) +
    scale_fill_manual(name = "",
                      labels = c(paste0("Soil Ca:Mg = ", ca_mg_min),
                                 paste0("Soil Ca:Mg = ", ca_mg_max)),
                      values = colpal) +
    scale_color_manual(name = "",
                       labels = c(paste0("Soil Ca:Mg = ", ca_mg_min),
                                  paste0("Soil Ca:Mg = ", ca_mg_max)),
                       values = colpal) +
    scale_y_log10() +
    xlab("log(Specific leaf area)") +
    ylab("Fecundity") +
    theme(legend.position = "top",
          legend.text = element_text(size = 12))


eff_sla_depth_df <- effects::effect("log_sla:depth", trait_model_full,
                                 xlevels = list(depth = c(min(perf_env_trt$depth),
                                                          max(perf_env_trt$depth)),
                                                log_sla = c(min(perf_env_trt$log_sla),
                                                            max(perf_env_trt$log_sla)))) %>%
  data.frame

plot_sla_depth <-
    ggplot(eff_sla_depth_df, aes(x = (log_sla*sla_sd)+sla_mean, y = fit, color=factor(depth)))+
    geom_line(size=1.25)+
    geom_ribbon(aes(ymin=lower, ymax=upper,fill=factor(depth)),
                alpha=0.3, colour= NA) +
    annotate("text", x = -Inf, y = Inf, hjust = 0, vjust = 1,
             label = get_pval(mod_term = "log_sla:depth")) +

    scale_fill_manual(name = "",
                      labels = c(paste0("Soil depth = ", depth_min, "cm"),
```

```
                                 paste0("Soil depth = ", depth_max, "cm")),
                      values = colpal) +
    scale_color_manual(name = "",
                       labels = c(paste0("Soil depth = ", depth_min, "cm"),
                                  paste0("Soil depth = ", depth_max, "cm")),
                       values = colpal) +
    scale_y_log10() +
    xlab("log(Specific leaf area)") +
    ylab("Fecundity") +
    theme(legend.position = "top",
          legend.text = element_text(size = 12))

eff_srl_sand_df <- effects::effect("sand:log_srl", trait_model_full,
                                xlevels = list(sand = c(min(perf_env_trt$sand),
                                                        max(perf_env_trt$sand)),
                                               log_srl = c(min(perf_env_trt$log_srl),
                                                           max(perf_env_trt$log_srl)))) %>%
  data.frame
plot_srl_sand <-
    ggplot(eff_srl_sand_df, aes(x = (log_srl*srl_sd) + srl_mean, y = fit, color=factor(sand)))+
    geom_line(size=1.25)+
    geom_ribbon(aes(ymin=lower,ymax=upper,fill=factor(sand)),
                alpha=0.3, colour= NA) +
    annotate("text", x = -Inf, y = Inf, hjust = 0, vjust = 1,
             label = get_pval(mod_term = "sand:log_srl")) +
    scale_fill_manual(name = "",
                      labels = c(paste0("Sand content = ", sand_min, "%"),
                                 paste0("Sand content = ", sand_max, "%")),
                      values = colpal) +
    scale_color_manual(name = "",
                       labels = c(paste0("Sand content = ", sand_min, "%"),
                                  paste0("Sand content = ", sand_max, "%")),
                       values = colpal) +
    scale_y_log10() +
    xlab("log(Specific root length)") +
    ylab("Fecundity") +
```

```r
    theme(legend.position = "top",
          legend.text = element_text(size = 12))

eff_maxht_depth_df <- effects::effect("depth:log_maxht", trait_model_full,
                                  xlevels = list(depth = c(min(perf_env_trt$depth),
                                                           max(perf_env_trt$depth)),
                                                 log_maxht = c(min(perf_env_trt$log_maxht),
                                                               max(perf_env_trt$log_maxht)))) %>%
  data.frame

plot_maxht_depth <-
    ggplot(eff_maxht_depth_df, aes(x = (log_maxht*maxht_sd) + maxht_mean, y = fit, color=factor(depth))
    geom_line(size=1.25)+
    geom_ribbon(aes(ymin=lower,ymax=upper,fill=factor(depth)),
                alpha=0.3, colour= NA) +
    annotate("text", x = -Inf, y = Inf, hjust = 0, vjust = 1,
             label = get_pval(mod_term = "depth:log_maxht")) +
    scale_fill_manual(name = "",
                      labels = c(paste0("Soil depth = ", depth_min, "cm"),
                                 paste0("Soil depth = ", depth_max, "cm")),
                      values = colpal) +
    scale_color_manual(name = "",
                       labels = c(paste0("Soil depth = ", depth_min, "cm"),
                                  paste0("Soil depth = ", depth_max, "cm")),
                       values = colpal) +
    scale_y_log10() +
    xlab("log(Maximum height)") +
    ylab("Fecundity") +
    theme(legend.position = "top",
          legend.text = element_text(size = 12))

lsla_camg_plot <- lsla_camg_plot + ylab(TeX("CWM log(SLA) (cm^2/mg)"))
lsla_depth_plot <- lsla_depth_plot + ylab(TeX("CWM log(SLA) (cm^2/mg)"))
lsrl_sand_plot <- lsrl_sand_plot + ylab("CWM log(SRL) (m/g)")
lmaxht_depth_plot <- lmaxht_depth_plot + ylab("CWM log(Max height) (cm)")
comboplots <-
```

```
  {lsla_camg_plot + lsla_depth_plot +
      lsrl_sand_plot + lmaxht_depth_plot +
      plot_sla_camg + plot_sla_depth +
      plot_srl_sand + plot_maxht_depth} +
  plot_layout(ncol = 2, byrow = F) +
  plot_annotation(tag_levels = "A")
comboplots
```

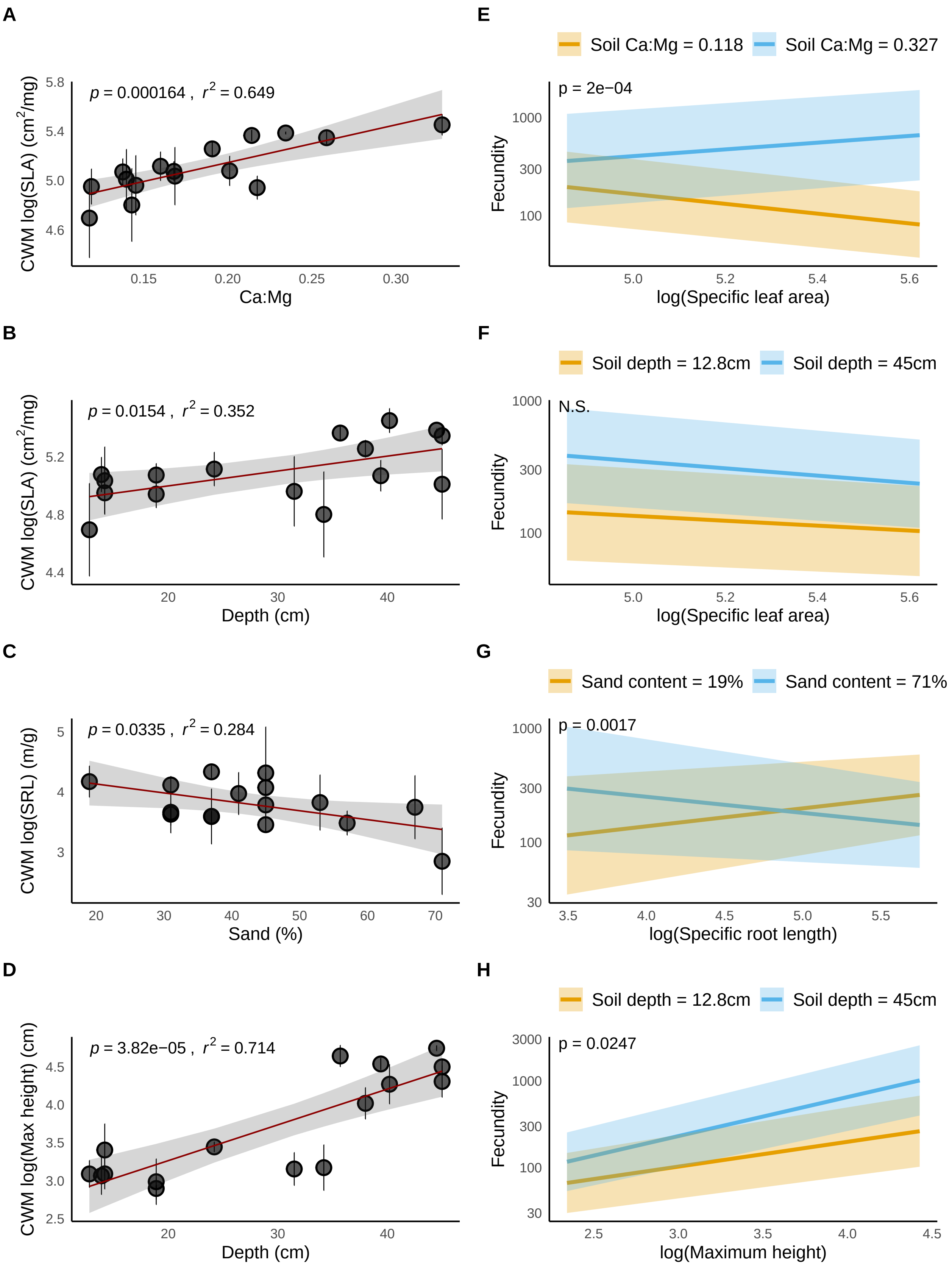
A
p = 0.000164 , r2 = 0.649
CWM log(SLA) (cm2/mg)
5.8
5.4
5.0
4.6
0.15
0.20
0.25
0.30
Ca:Mg
E
Soil Ca:Mg = 0.118
Soil Ca:Mg = 0.327
p = 2e−04
Fecundity
1000
300
100
5.0
5.2
5.4
5.6
log(Specific leaf area)
B
p = 0.0154 , r2 = 0.352
CWM log(SLA) (cm2/mg)
5.2
4.8
4.4
20
30
40
Depth (cm)
F
Soil depth = 12.8cm
Soil depth = 45cm
N.S.
Fecundity
1000
300
100
5.0
5.2
5.4
5.6
log(Specific leaf area)
C
p = 0.0335 , r2 = 0.284
CWM log(SRL) (m/g)
5
4
3
20
30
40
50
60
70
Sand (%)
G
Sand content = 19%
Sand content = 71%
p = 0.0017
Fecundity
1000
300
100
30
3.5
4.0
4.5
5.0
5.5
log(Specific root length)
D
p = 3.82e−05 , r2 = 0.714
CWM log(Max height) (cm)
4.5
4.0
3.5
3.0
2.5
20
30
40
Depth (cm)
H
Soil depth = 12.8cm
Soil depth = 45cm
p = 0.0247
Fecundity
3000
1000
300
100
30
2.5
3.0
3.5
4.0
4.5
log(Maximum height)

```r
ggsave("manuscript/figures/cwm-tXe-plots.pdf", height = 12, width = 9)
```

189 Visualize interactions surfaces

```r
# Visualize a 3D-surface
# pdf("manuscript/figures/supplemental/tXe-surface.pdf", width = 9, height = 9)
par(mfrow = c(3,3))
visreg2d(trait_model_full, "log_sla", "ca_mg", plot.type = "persp", theta = 30,
         xlab = "SLA (Scaled)", ylab = "Ca:Mg (Scaled)", zlab = "Fecundity",
         scale = "response", main = "SLA x Ca:Mg*")
visreg2d(trait_model_full, "log_sla", "sand", plot.type = "persp", theta = 30,
         xlab = "SLA (Scaled)", ylab = "Soil sand content (Scaled)", zlab = "Fecundity",
         scale = "response", main = "SLA x Sand content")
visreg2d(trait_model_full, "log_sla", "depth", plot.type = "persp", theta = 30,
         xlab = "SLA (Scaled)", ylab = "Soil depth (Scaled)", zlab = "Fecundity",
         scale = "response", main = "SLA x Soil depth*")
visreg2d(trait_model_full, "log_srl", "ca_mg", plot.type = "persp", theta = 30,
         xlab = "SRL (Scaled)", ylab = "Ca:Mg (Scaled)", zlab = "Fecundity",
         scale = "response", main = "SRL x Ca:Mg")
visreg2d(trait_model_full, "log_srl", "sand", plot.type = "persp", theta = 40,
         xlab = "SRL (Scaled)", ylab = "Sand (Scaled)", zlab = "Fecundity",
         scale = "response", main = "SRL x Sand content*")
visreg2d(trait_model_full, "log_srl", "depth", plot.type = "persp", theta = 40,
         xlab = "SRL (Scaled)", ylab = "Soil depth (Scaled)", zlab = "Fecundity",
         scale = "response", main = "SRL x Soil depth")
visreg2d(trait_model_full, "log_maxht", "ca_mg", plot.type = "persp", theta = 30,
         xlab = "Max Height (Scaled)", ylab = "Ca:Mg (Scaled)", zlab = "Fecundity",
         scale = "response", main = "Max height x Ca:Mg depth")
visreg2d(trait_model_full, "log_maxht", "sand", plot.type = "persp", theta = 30,
         xlab = "Max Height (Scaled)", ylab = "Sand content (Scaled)", zlab = "Fecundity",
         scale = "response", main = "Max height x Sand content")
visreg2d(trait_model_full, "log_maxht", "depth", plot.type = "persp", theta = 30,
         xlab = "Max Height (Scaled)", ylab = "Depth (Scaled)", zlab = "Fecundity",
         scale = "response", main = "Max height x Soil depth*")
```

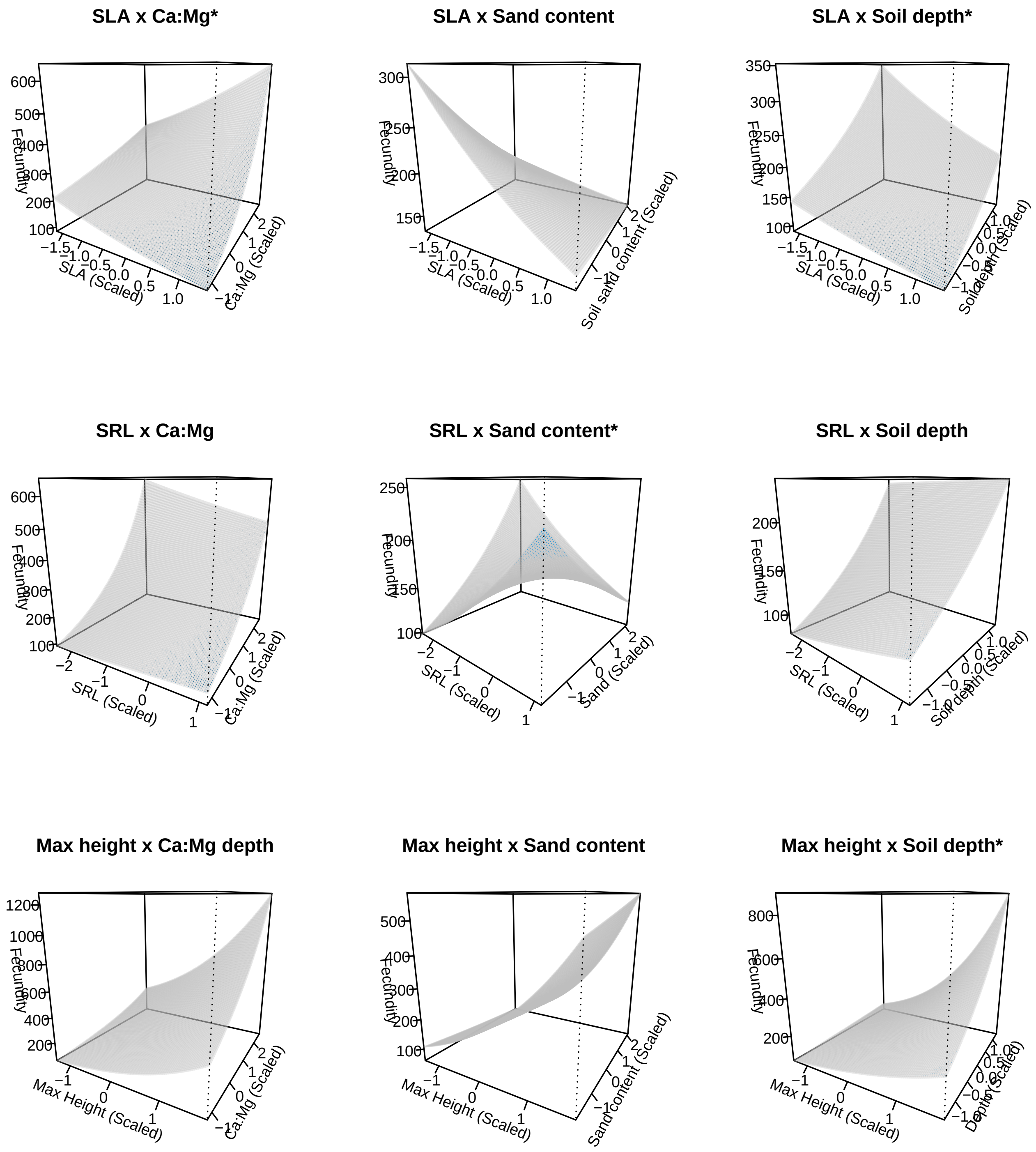


```
# dev.off()
```

Now, make plots of all the non-significant T-E interactions:

```
# Make plots of all non-sig tXe ----


eff_sla_sand_df <- effects::effect("log_sla:sand", trait_model_full,
```

```
                                  xlevels = list(sand = c(min(perf_env_trt$sand),
                                                         max(perf_env_trt$sand)),
                                               log_sla = c(min(perf_env_trt$log_sla),
                                                             max(perf_env_trt$log_sla)))) %>%
  data.frame
plot_sla_sand <-
    ggplot(eff_sla_sand_df, aes(x = (log_sla*sla_sd)+sla_mean, y = fit, color=factor(sand)))+
    geom_line(size=1.25)+
    geom_ribbon(aes(ymin=lower,ymax=upper, fill=factor(sand)),
                alpha=0.3, colour= NA) +
    annotate("text", x = -Inf, y = Inf, hjust = 0, vjust = 1,
             label = get_pval(mod_term = "log_sla:sand")) +
    scale_fill_manual(name = "",
                      labels = c(paste0("Soil sand = ", sand_min),
                                 paste0("Soil sand = ", sand_max)),
                      values = colpal) +
    scale_color_manual(name = "",
                       labels = c(paste0("Soil sand = ", sand_min),
                                  paste0("Soil sand = ", sand_max)),
                       values = colpal) +
    scale_y_log10() +
    xlab("log(Specific leaf area)") +
    ylab("Fecundity") +
    theme(legend.position = "top",
          legend.text = element_text(size = 12))


eff_srl_camg_df <- effects::effect("ca_mg:log_srl", trait_model_full,
                                  xlevels = list(ca_mg = c(min(perf_env_trt$ca_mg),
                                                          max(perf_env_trt$ca_mg)),
                                               log_srl = c(min(perf_env_trt$log_srl),
                                                             max(perf_env_trt$log_srl)))) %>%
  data.frame
plot_srl_camg <-
    ggplot(eff_srl_camg_df, aes(x = (log_srl*srl_sd)+sla_mean, y = fit, color=factor(ca_mg)))+
    geom_line(size=1.25)+
```

```
    geom_ribbon(aes(ymin=lower,ymax=upper, fill=factor(ca_mg)),
                alpha=0.3, colour= NA) +
    annotate("text", x = -Inf, y = Inf, hjust = 0, vjust = 1,
             label = get_pval(mod_term = "ca_mg:log_srl")) +
    scale_fill_manual(name = "",
                      labels = c(paste0("Soil Ca:Mg = ", ca_mg_min),
                                 paste0("Soil Ca:Mg = ", ca_mg_max)),
                      values = colpal) +
    scale_color_manual(name = "",
                       labels = c(paste0("Soil Ca:Mg = ", ca_mg_min),
                                  paste0("Soil Ca:Mg = ", ca_mg_max)),
                       values = colpal) +
    scale_y_log10() +
    xlab("log(Specific root length)") +
    ylab("Fecundity") +
    theme(legend.position = "top",
          legend.text = element_text(size = 12))


eff_srl_depth_df <- effects::effect("depth:log_srl", trait_model_full,
                                    xlevels = list(depth = c(min(perf_env_trt$depth),
                                                             max(perf_env_trt$depth)),
                                                   log_srl = c(min(perf_env_trt$log_srl),
                                                               max(perf_env_trt$log_srl)))) %>%
  data.frame
plot_srl_depth <-
    ggplot(eff_srl_depth_df, aes(x = (log_srl*srl_sd)+sla_mean, y = fit, color=factor(depth)))+
    geom_line(size=1.25)+
    geom_ribbon(aes(ymin=lower,ymax=upper, fill=factor(depth)),
                alpha=0.3, colour= NA) +
    annotate("text", x = -Inf, y = Inf, hjust = 0, vjust = 1,
             label = get_pval(mod_term = "depth:log_srl")) +
    scale_fill_manual(name = "",
                      labels = c(paste0("Soil Depth = ", depth_min),
                                 paste0("Soil Depth = ", depth_max)),
                      values = colpal) +
```

```
    scale_color_manual(name = "",
                       labels = c(paste0("Soil Depth = ", depth_min),
                                  paste0("Soil Depth = ", depth_max)),
                       values = colpal) +
    scale_y_log10() +
    xlab("log(Specific root length)") +
    ylab("Fecundity") +
    theme(legend.position = "top",
          legend.text = element_text(size = 12))
```

```
eff_maxht_sand_df <- effects::effect("sand:log_maxht", trait_model_full,
                                     xlevels = list(sand = c(min(perf_env_trt$sand),
                                                             max(perf_env_trt$sand)),
                                                    log_maxht = c(min(perf_env_trt$log_maxht),
                                                                  max(perf_env_trt$log_maxht)))) %>%
  data.frame
plot_maxht_sand <-
    ggplot(eff_maxht_sand_df, aes(x = (log_maxht*maxht_sd)+maxht_mean, y = fit, color=factor(sand)))+
    geom_line(size=1.25)+
    geom_ribbon(aes(ymin=lower,ymax=upper, fill=factor(sand)),
                alpha=0.3, colour= NA) +
    annotate("text", x = -Inf, y = Inf, hjust = 0, vjust = 1,
             label = get_pval(mod_term = "sand:log_maxht")) +
    scale_fill_manual(name = "",
                      labels = c(paste0("Soil sand = ", sand_min),
                                 paste0("Soil sand = ", sand_max)),
                      values = colpal) +
    scale_color_manual(name = "",
                       labels = c(paste0("Soil sand = ", sand_min),
                                  paste0("Soil sand = ", sand_max)),
                       values = colpal) +
    scale_y_log10() +
```

```
    xlab("log(Max height)") +
    ylab("Fecundity") +
    theme(legend.position = "top",
          legend.text = element_text(size = 12))



eff_maxht_camg_df <- effects::effect("ca_mg:log_maxht", trait_model_full,
                                     xlevels = list(ca_mg = c(min(perf_env_trt$ca_mg),
                                                              max(perf_env_trt$ca_mg)),
                                                    log_maxht = c(min(perf_env_trt$log_maxht),
                                                                  max(perf_env_trt$log_maxht)))) %>%
  data.frame
plot_maxht_camg <-
    ggplot(eff_maxht_camg_df, aes(x = (log_maxht*maxht_sd)+maxht_mean, y = fit, color=factor(ca_mg)))+
    geom_line(size=1.25)+
    geom_ribbon(aes(ymin=lower,ymax=upper, fill=factor(ca_mg)),
                alpha=0.3, colour= NA) +
    annotate("text", x = -Inf, y = Inf, hjust = 0, vjust = 1,
             label = get_pval(mod_term = "ca_mg:log_maxht")) +
    scale_fill_manual(name = "",
                      labels = c(paste0("Soil Ca:Mg = ", ca_mg_min),
                                 paste0("Soil Ca:Mg = ", ca_mg_max)),
                      values = colpal) +
    scale_color_manual(name = "",
                       labels = c(paste0("Soil Ca:Mg = ", ca_mg_min),
                                  paste0("Soil Ca:Mg = ", ca_mg_max)),
                       values = colpal) +
    scale_y_log10() +
    xlab("log(Max height)") +
    ylab("Fecundity") +
    theme(legend.position = "top",
          legend.text = element_text(size = 12))

comboplots_supplemental <-
  {lsrl_camg_plot + lsrl_depth_plot + lsla_sand_plot +
      lmaxht_sand_plot + lmaxht_camg_plot + plot_srl_camg +
```

```
      plot_srl_depth + plot_sla_sand + plot_maxht_sand +
      plot_maxht_camg} +
  plot_layout(ncol = 2, byrow = F) +
  plot_annotation(tag_levels = "A")
ggsave("manuscript/figures/supplemental/cwm-tXe-plots-supplemental.pdf",
       comboplots_supplemental, height = 12, width = 9)

comboplots_supplemental
```

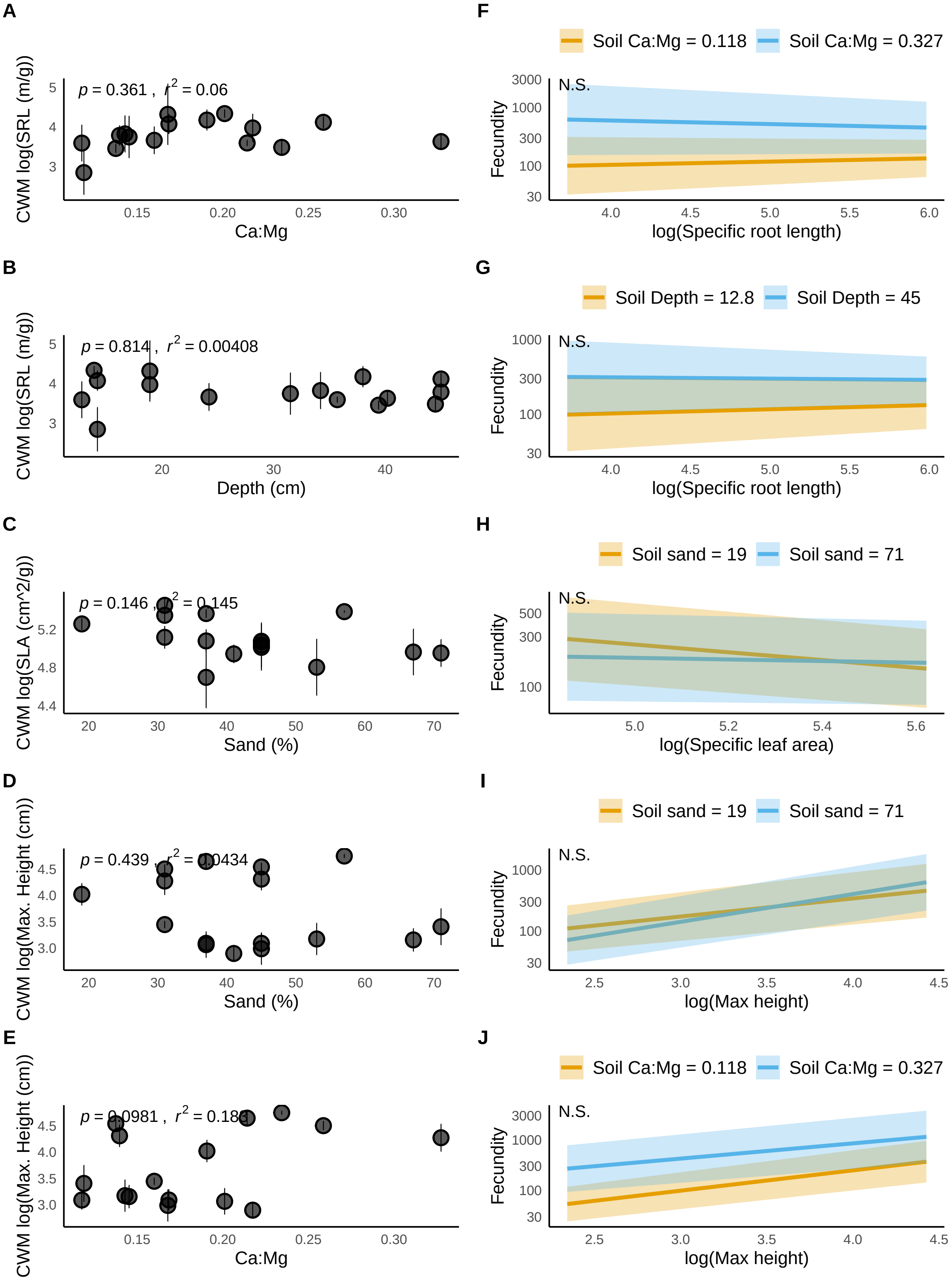
A
p = 0.361 , r² = 0.06
CWM log(SRL (m/g))
Ca:Mg
B
p = 0.814 , r² = 0.00408
CWM log(SRL (m/g))
Depth (cm)
C
p = 0.146 , r² = 0.145
CWM log(SLA (cm^2/g))
Sand (%)
D
p = 0.439 , r² = 0.0434
CWM log(Max. Height (cm))
Sand (%)
E
p = 0.0981 , r² = 0.183
CWM log(Max. Height (cm))
Ca:Mg
F
Soil Ca:Mg = 0.118
Soil Ca:Mg = 0.327
N.S.
Fecundity
log(Specific root length)
G
Soil Depth = 12.8
Soil Depth = 45
N.S.
Fecundity
log(Specific root length)
H
Soil sand = 19
Soil sand = 71
N.S.
Fecundity
log(Specific leaf area)
I
Soil sand = 19
Soil sand = 71
N.S.
Fecundity
log(Max height)
J
Soil Ca:Mg = 0.118
Soil Ca:Mg = 0.327
N.S.
Fecundity
log(Max height)

### Supplemental analyses

#### Quadratic CWM trait-env models

```
p17_sitesum_env$ca_mg2 <- (p17_sitesum_env$ca_mg)^2
p17_sitesum_env$depth2 <- (p17_sitesum_env$depth)^2
p17_sitesum_env$sand2 <- (p17_sitesum_env$sand)^2

lsla_camg2_lm <- lm(lsla_cwm_mean~ca_mg + ca_mg2, data = p17_sitesum_env)
lsla_depth2_lm <- lm(lsla_cwm_mean~depth + depth2, data = p17_sitesum_env)
lsla_sand2_lm <- lm(lsla_cwm_mean~sand + sand2, data = p17_sitesum_env)

lsrl_camg2_lm <- lm(lsrl_cwm_mean~ca_mg + ca_mg2, data = p17_sitesum_env)
lsrl_depth2_lm <- lm(lsrl_cwm_mean~depth + depth2, data = p17_sitesum_env)
lsrl_sand2_lm <- lm(lsrl_cwm_mean~sand + sand2, data = p17_sitesum_env)

lmaxht_camg2_lm <- lm(lmaxht_cwm_mean~ca_mg + ca_mg2, data = p17_sitesum_env)
lmaxht_depth2_lm <- lm(lmaxht_cwm_mean~depth + depth2, data = p17_sitesum_env)
lmaxht_sand2_lm <- lm(lmaxht_cwm_mean~sand + sand2, data = p17_sitesum_env)

quadmodels_list <- list(lsla_camg2_lm, lsla_depth2_lm, lsla_sand2_lm,
                        lsrl_camg2_lm, lsrl_depth2_lm, lsrl_sand2_lm,
                        lmaxht_camg2_lm, lmaxht_depth2_lm, lmaxht_sand2_lm)

quadmodels_df <- map_df(quadmodels_list, tidy)

quadmodels_df$trait <- c("SLA", rep("", 8),
                         "SRL", rep("", 8),
                         "Max. Height", rep("", 8))
quadmodels_df$trait <- rep(c("SLA", "SRL", "Max. Height"), each = 9)
quadmodels_df$environment <- rep(c("Ca:Mg" ,"","",
                                   "Depth", "", "",
                                   "Sand", "", ""), 3)
quadmodels_df$term <- rep(c("Intercept", "Linear", "Quadratic"), 9)
quadmodels_df$std.error <- round(quadmodels_df$std.error, 4)
quadmodels_df$statistic <- round(quadmodels_df$statistic, 4)
quadmodels_df$p.value <- round(quadmodels_df$p.value, 4)
```

```r
quadmodels_df <- quadmodels_df %>% select(trait, environment, term, estimate, std.error, statistic,
                                          p.value)

quadmodels_df_print <- knitr::kable(quadmodels_df, format = "latex", booktabs = T,
                                    linesep = c(rep(c("", "",  "\\addlinespace",
                                                      "", "",  "\\addlinespace",
                                                      "", "",  "\\addlinespace \\hline"), 3)),
                                    caption = "Model output for quadratic relationships between CWM trai
                                    caption.short = "Testing for quadratic relationships between CWM tra
                                    label = "CWMquadratic") %>%
  kableExtra::kable_styling()
saveRDS(object = quadmodels_df_print, file = "manuscript/figures/quadmodels_df.Rds")
```

#### Environment PCA

```r
environment <- read_csv("data/environmental/all_environmental_data.csv") %>%
  rename(plot_num = plot) %>%
  mutate(microsite = ifelse(plot_num < 756, microsite, "lower")) %>%
  filter(microsite != "lower") %>%
  return
```

```
## Parsed with column specification:
## cols(
##   .default = col_double(),
##   type = col_character(),
##   microsite = col_character()
## )

## See spec(...) for full column specifications.
```

```r
colnames(environment)
```

```
##  [1] "plot_num"           "lat"                "lon"
##  [4] "ele"                "type"               "Tmax"
##  [7] "Tmin"               "organic_matter_ENR" "pH"
## [10] "CEC_meq_100g"       "K_ppm"              "Mg_ppm"
## [13] "Ca_ppm"             "NH4_N_ppm"          "Nitrate_ppm"
## [16] "soil_moisture"      "sand"               "clay"
```

```
209 ## [19] "microsite"          "depth"              "P1_weak_bray_ppm"
210 ## [22] "NaHCO3P_olsen_ppm"
```

```
environment$ca_mg <- environment$Ca_ppm/environment$Mg_ppm


environment_pcacols <- environment %>%
  select(plot_num, organic_matter_ENR, pH, CEC_meq_100g,
         K_ppm, Ca_ppm, Mg_ppm, NH4_N_ppm, Nitrate_ppm,
         soil_moisture, sand, depth, Tmax, ca_mg,  P1_weak_bray_ppm) %>%
  mutate(plot_num = as.character(plot_num)) %>%
  # scale the columns
  mutate_if(is.numeric, scale)  %>%
  # plot number should be the name of the rows.
  column_to_rownames("plot_num")
env_pca <- PCA(environment_pcacols, graph = F)


env_pca_coords <- env_pca$ind$coord
env_pca_coords <- env_pca_coords %>% data.frame %>% rownames_to_column("plot_num") %>%
  mutate(plot_num = as.numeric(plot_num)) %>%
  left_join(., environment %>% select(plot_num, microsite))
```

```
211 ## Joining, by = "plot_num"
```

```
env_pca_12 <- factoextra::fviz_pca_biplot(env_pca, col.ind = "transparent") +
  geom_point(data = env_pca_coords, aes(x = Dim.1, y = Dim.2, shape = microsite), size = 4)+
  scale_shape_manual(values = c(20,21), labels = c("hummock", "matrix")) +
  theme(legend.position = "bottom")+
  labs(tile = "")

env_pca_23 <- factoextra::fviz_pca_biplot(env_pca, axes = c(2,3), col.ind = "transparent")+
  geom_point(data = env_pca_coords, aes(x = Dim.2, y = Dim.3, shape = microsite), size = 4)+
  scale_shape_manual(values = c(20,21), labels = c("hummock", "matrix")) +
  theme(legend.position = "bottom") +
  labs(tile = "")



env_pcas_figure <- {env_pca_12 + env_pca_23} +
  plot_annotation(tag_levels = "A") +
```

```
  plot_layout(guides = 'collect') &
  theme(legend.position = "bottom")
env_pcas_figure
```

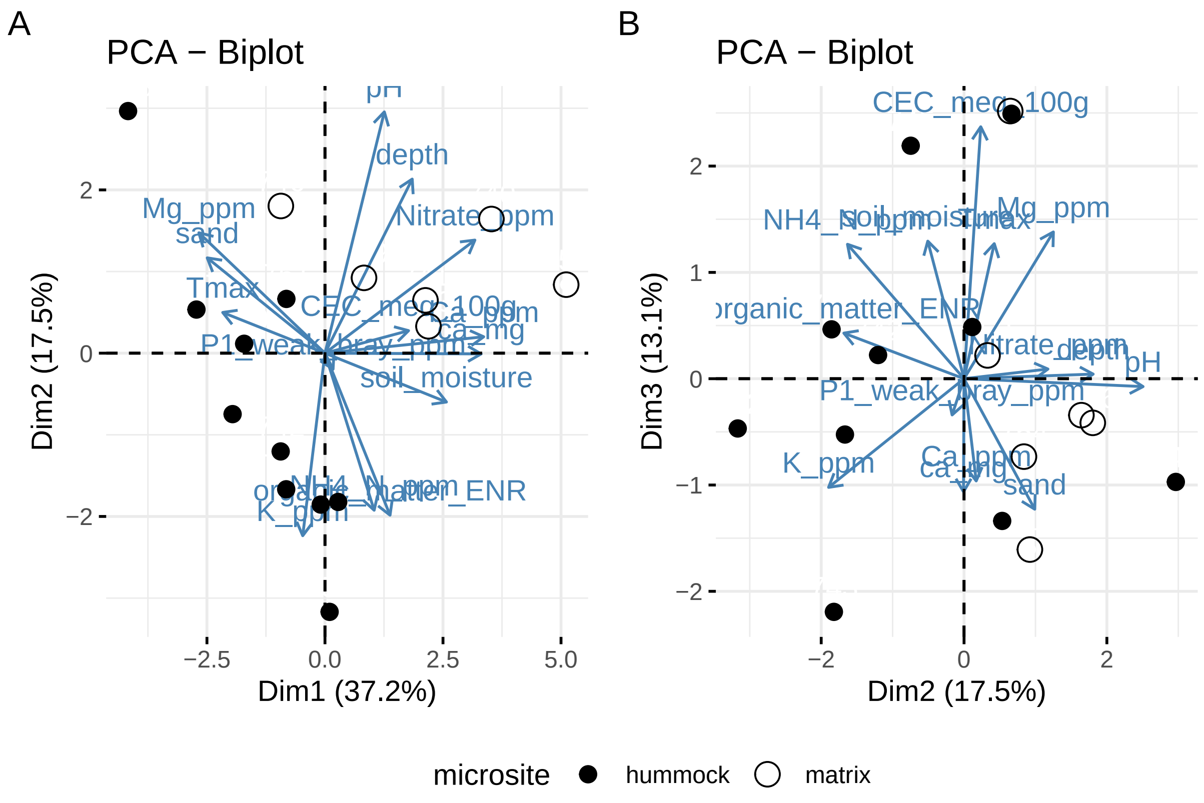


```
ggsave("manuscript/figures/supplemental/env-pca.pdf", height = 6, width = 12)
```

##### Trait PCA

```
na_count <- sapply(strts, function(y) sum(length(which(is.na(y)))))
na_count
```

```
##          USDA_symbol           leaf_size                 SLA
##                    0                   1                   1
##                 LDMC                 LAI                 LAR
##                    1                  10                   7
##            seed_mass          max_height                 SRL
##                    1                   0                  16
##      relative_spread      projected_area           phenology
##                   12                  10                  22
##            foliar_N            CN_ratio                d13C
```

```
##                       7                       7                       7
##                    d15N                   notes   seed_mass_data_source
##                       7                      54                       1
## max_height_data_source                 dataset       turgor_loss_point
##                       0                       0                      38
##               seed_size           rooting_depth                 leaf_pH
##                      40                      40                      40
```

```r
# Exclude traits for which we are lacking data from most
# species (leaf pH, TLP, rooting depth, seed 'size)
# also exclude notes/non-numeric columns for PCA
strts <- strts %>% select(-c(leaf_pH, turgor_loss_point, dataset,
                             max_height_data_source, LAI, LAR,
                             seed_mass_data_source, notes, rooting_depth,
                             seed_size, projected_area))
# Species codes for the focal species of the experiment
focal_sp_codes <- c("URLI5", "SACO6", "PLER3", "MICA",
                    "MEPO3", "LOWR2", "LACA7", "HOMU",
                    "HECO7","FEMI2", 'EUSP', "CLPU2",
                    'CLBO', 'CHGL', 'CEME2', 'BRMA3', 'AMME')
dominant_sp_codes <- c("AVFA", "BRDI3", "AVBA", "MIDO", "LOPE")
focal_sp_rownums <- which(strts$USDA_symbol %in% focal_sp_codes)

# Scale the numeric trait columns
strts_scaled <- strts %>%
  column_to_rownames("USDA_symbol") %>%
  replace_na(.,as.list(colMeans(.,na.rm=T))) %>%
  mutate_if(is.numeric, scale)

trait_pca <- PCA(strts_scaled, graph = F)
focal_sp_coords <- data.frame(trait_pca$ind$coord[focal_sp_rownums,])

focal_sp_codes_sedgnames <- c("URLI", "SACO", "PLER", "MICA",
                              "MEPO", "ACWR", 'LACA', 'HOMU',
                              'HECO', 'VUMI', 'EUSP', 'CLPU',
                              'CLBO', 'CHGL', 'CEME', 'BRMA', 'AMME')
focal_sp_coords$sp_code <- focal_sp_codes_sedgnames
```

```r
trait_pca_12 <- fviz_pca_biplot(trait_pca, col.ind = "grey", label = "var") +
  labs(title = "") +
  geom_point(data = focal_sp_coords, aes(x = Dim.1, y = Dim.2),
             color = "darkred", size = 2.5) +
  ggrepel::geom_text_repel(data = focal_sp_coords, aes(x = Dim.1, y = Dim.2, label = sp_code))

trait_pca_23 <- fviz_pca_biplot(trait_pca, col.ind = "grey", label = "var", axes = c(2,3)) +
  labs(title = "") +
  geom_point(data = focal_sp_coords, aes(x = Dim.2, y = Dim.3),
             color = "darkred", size = 2.5) +
  ggrepel::geom_text_repel(data = focal_sp_coords, aes(x = Dim.2, y = Dim.3, label = sp_code))

trait_pca_plot <- {trait_pca_12 + trait_pca_23} +
  patchwork::plot_annotation(tag_levels = "A")
trait_pca_plot
```

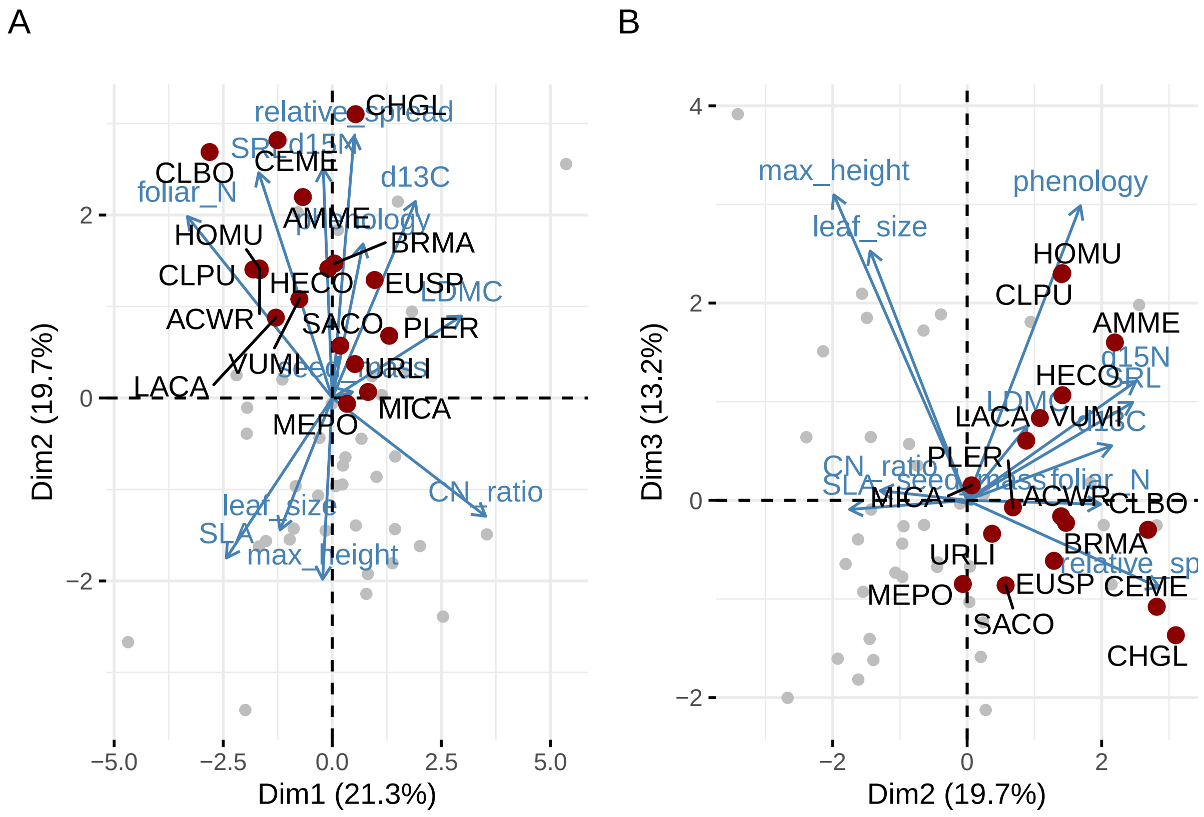


```r
saveRDS(trait_pca_plot, "manuscript/figures/supplemental/trait-pca.Rds")
ggsave("manuscript/figures/supplemental/trait-pca.pdf",
       trait_pca_plot, height = 6, width = 12)
```

```r
maxht_hist <- ggplot(strts) +
  geom_histogram(aes(x = max_height), bins = 20) +
  geom_rug(aes(x = max_height), color = "#E69F00", size = 2, alpha = .75,
           data = strts %>% filter(USDA_symbol %in% focal_sp_codes)) +
  geom_rug(aes(x = max_height), color = "#0072B2", size = 2, alpha = .75,
           data = strts %>% filter(USDA_symbol %in% dominant_sp_codes)) +
  scale_x_log10() +
  xlab("Maximum height (cm)")

sla_hist <- ggplot(strts) +
  geom_histogram(aes(x = SLA), bins = 20) +
  geom_rug(aes(x = SLA), color = "#E69F00", size = 2, alpha = .75,
           data = strts %>% filter(USDA_symbol %in% focal_sp_codes)) +
  geom_rug(aes(x = SLA), color = "#0072B2", size = 2, alpha = .75,
           data = strts %>% filter(USDA_symbol %in% dominant_sp_codes)) +
  scale_x_log10() +
  xlab(TeX("Specific leaf area (cm^2/g)"))


srl_hist <- ggplot(strts) +
  geom_histogram(aes(x = SRL), bins = 20) +
  geom_rug(aes(x = SRL), color = "#E69F00", size = 2, alpha = .75,
           data = strts %>% filter(USDA_symbol %in% focal_sp_codes)) +
  geom_rug(aes(x = SRL), color = "#0072B2", size = 2, alpha = .75,
           data = strts %>% filter(USDA_symbol %in% dominant_sp_codes)) +
  scale_x_log10() +
  xlab("Specific root length (m/g)")

fig_traithistograms <- {maxht_hist + sla_hist + srl_hist} +
  plot_annotation(tag_levels = "A")
fig_traithistograms
```

```
## Warning: Removed 1 rows containing non-finite values (stat_bin).

## Warning: Removed 16 rows containing non-finite values (stat_bin).
```

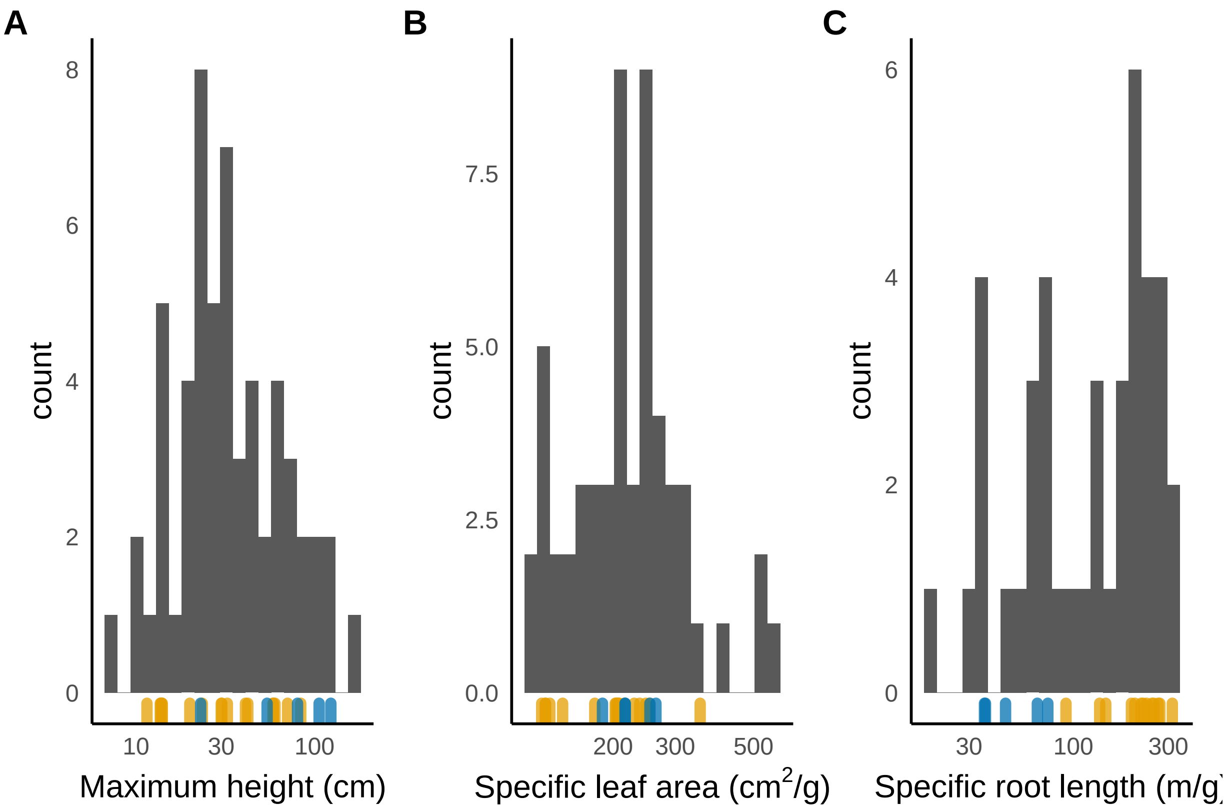


```r
ggsave("manuscript/figures/supplemental/trait-histograms.pdf",
       fig_traithistograms, width = 12, height = 4)
```

```
## Warning: Removed 1 rows containing non-finite values (stat_bin).

## Warning: Removed 16 rows containing non-finite values (stat_bin).
```

```r
trait_table <- data.frame(
  Organ = c("Whole plant"," "," "," ", " ", "Leaf", "", "", "", "", "Root"),
  Trait = c("Max. height", "Canopy shape index", "Carbon isotope composition (dC13)",
            "Phenology", "Seed mass", "Leaf size", "Specific leaf area",
            "Leaf dry matter content", "C:N ratio", "Leaf N concentration",
            "Specific root length"),
  Units = c("cm", "dimensionless", "dC13", "day of year", "mg", "cm$^2$",
            "g/cm$^2$", "mg/g", "dimensionless", "mg/g", "m/g")
)
trait_table_print <- knitr::kable(trait_table, format = "latex", booktabs = T,
                                  label = "traittable",
                                  caption = "List of traits measured for this study") %>%
  kableExtra::kable_styling()
```

```r
saveRDS(object = trait_table_print, file = "manuscript/figures/trait_table.Rds")
```
